# The role of redox-cofactor regeneration and ammonium assimilation in secretion of amino acids as byproducts of *Clostridium thermocellum*

**DOI:** 10.1101/2022.10.12.512009

**Authors:** Johannes Yayo, Thomas Rydzak, Teun Kuil, Anna Karlsson, Dan J. Harding, Adam M. Guss, Antonius J. A. van Maris

## Abstract

*Clostridium thermocellum* is a cellulolytic thermophile considered for consolidated bioprocessing of lignocellulose to ethanol. Improvements in ethanol yield are required for industrial implementation, but incompletely understood causes of amino acid secretion impede progress. In this study, amino acid secretion was investigated by gene deletions in ammonium-regulated NADPH-supplying and -consuming pathways and physiological characterization in cellobiose- or ammonium-limited chemostats. First, the contribution of the NADPH-supplying malate shunt was studied with strains using either the NADPH-yielding malate shunt (Δ*ppdk*) or redox-independent conversion of PEP to pyruvate (Δ*ppdk* Δ*malE::P_eno_-pyk*). In the latter, branched-chain amino acids, especially valine, were significantly reduced, whereas the ethanol yield increased 46-60%, suggesting that secretion of these amino acids balances NADPH surplus from the malate shunt. Unchanged amino acid secretion in Δ*ppdk* falsified a previous hypothesis on ammonium-regulated PEP-to-pyruvate flux redistribution. Possible involvement of another NADPH-supplier, namely NADH-dependent reduced ferredoxin:NADP^+^ oxidoreductase (*nfnAB*), was also excluded. Finally, deletion of glutamate synthase (*gogat*) in ammonium assimilation resulted in upregulation of NADPH-linked glutamate dehydrogenase activity and decreased amino acid yields. Since *gogat* in *C. thermocellum* is putatively annotated as ferredoxin-linked, which is supported by product redistribution observed in this study, this deletion likely replaced ferredoxin with NADPH in ammonium assimilation. Overall, these findings indicate that a need to reoxidize NADPH is driving the observed amino acid secretion, likely at the expense of NADH needed for ethanol formation. This suggests that metabolic engineering strategies on simplifying redox metabolism and ammonium assimilation can contribute to increased ethanol yields.

**Importance:** Improving the ethanol yield of *C. thermocellum* is important for industrial implementation of this microorganism in consolidated bioprocessing. A central role of NADPH in driving amino acid byproduct formation was demonstrated, by eliminating the NADPH-supplying malate shunt and separately by changing the cofactor specificity in ammonium assimilation. With amino acid secretion diverting carbon and electrons away from ethanol, these insights are important for further metabolic engineering to reach industrial requirements on ethanol yield. This study also provides chemostat data relevant for training genome-scale metabolic models and improving the validity of their predictions, especially considering the reduced degree-of-freedom in redox metabolism of the strains generated here. In addition, this study advances fundamental understanding on mechanisms underlying amino acid secretion in cellulolytic Clostridia as well as regulation and cofactor specificity in ammonium assimilation. Together, these efforts aid development of *C. thermocellum* for sustainable consolidated bioprocessing of lignocellulose to ethanol with minimal pretreatment.

## INTRODUCTION

*Clostridium thermocellum* is a promising candidate for cost-competitive cellulosic ethanol production through consolidated bioprocessing due to its native ability to efficiently solubilize lignocellulose (1). This anaerobic thermophile (also called *Ruminiclostridium thermocellum*, *Hungateiclostridium thermocellum*, and *Acetivibrio thermocellus* (2)) outperforms other cellulosic microorganisms as well as fungal cellulases in lignocellulose solubilization (3). Recently, this microorganism has been considered for hybrid biological/catalytic conversion of cellulosic biomass to fuels, where ethanol produced by *C. thermocellum* is chemically upgraded to larger fuel molecules compatible with heavy-duty, difficult-to-electrify transportation modes (4). However, industrial implementation of *C. thermocellum* would require improvements of the hitherto achieved ethanol titer (30 g L^-1^ (5)) and yield (75% of theoretical maximum (6)) to >40 g L^-1^ at >90% of theoretical maximum yield (4). Increased understanding of the complex and flexible central metabolism of *C. thermocellum* would directly benefit the design-build-test-learn cycles aiming to reach these targets.

Secretion of amino acids as byproducts is an atypical phenomenon observed in both wild-type and engineered *C. thermocellum* strains that diverts sugar away from ethanol formation. Amino acid secretion occurs at both low (5 g L^-1^) and high (93 g L^-1^) loadings of cellulose in batch cultures, where up to 8.7% of the total consumed carbon is secreted as amino acids, mainly valine (up to 6.0% of total carbon) and alanine (up to 1.6% of total carbon) (7–9). In strains engineered and evolved for high ethanol titer (22 g L^-1^) and yield (75% of theoretical maximum) on cellulose, amino acid secretion increases to 10% of the consumed carbon, becoming the second most abundant organic product after ethanol (6). Redirecting this carbon, and the associated available electrons, from amino acids to ethanol, will, especially in view of the low profit margins for production of commodity chemicals and fuels, have a large impact on the economic viability of *C. thermocellum* based ethanol production.

Several hypotheses have been proposed for this unexpected amino acid secretion. In one hypothesis, the high nitrogen content in batch medium is thought to result in accumulation of amino acids (8). Based on this, it was expected that amino acid secretion would be lower in nitrogen-limited conditions. Surprisingly, a study by Holwerda et al. (10) showed the opposite effect, where nitrogen-limited chemostats resulted in 6-fold higher production rate of secreted amino acids than carbon-limited chemostats. Specifically, valine was most abundant, with a 20-fold increase, followed by an 8-fold increase in isoleucine, and a coinciding 50-fold higher pyruvate secretion. Cell lysis was excluded as a major mechanism based on calculations on the amino acid distribution in the protein fraction of biomass and observed concentrations and patterns in extracellular amino acids (10). The observation of increased amino acid formation under nitrogen limitation at otherwise identical growth rates in chemostats also makes it unlikely that increased reversibility of the tRNA charging in protein synthesis, caused by *C. thermocellum* lacking a cytosolic pyrophosphatase, is a major contributor to amino acid secretion (10). A third hypothesis by van der Veen et al. (7) proposes that amino acid secretion in *C. thermocellum* is related to redox-cofactor balancing. Investigating this hypothesis requires understanding of sources and sinks of especially NADPH and the regulation of the involved pathways by, amongst others, ammonium. Underlying the hypothesis of van der Veen et al. (7) is the reoxidation of NADPH in amino acid biosynthesis. Based on a current omics-based genome-scale metabolic model (GEM) of *C. thermocellum* (11), and extrapolation from model microorganisms, biosynthetic pathways of amino acids from their central building block are expected to predominantly use NADPH (Table 1). For instance, production of valine from two pyruvate requires two NADPH (Table 1). When considering this possible role of amino acid secretion as an NADPH sink, it is important to note that, in *C. thermocellum*, NADPH can additionally be reoxidized through hydrogen formation (Fig. 1C) (12, 13).

**FIG. 1.**
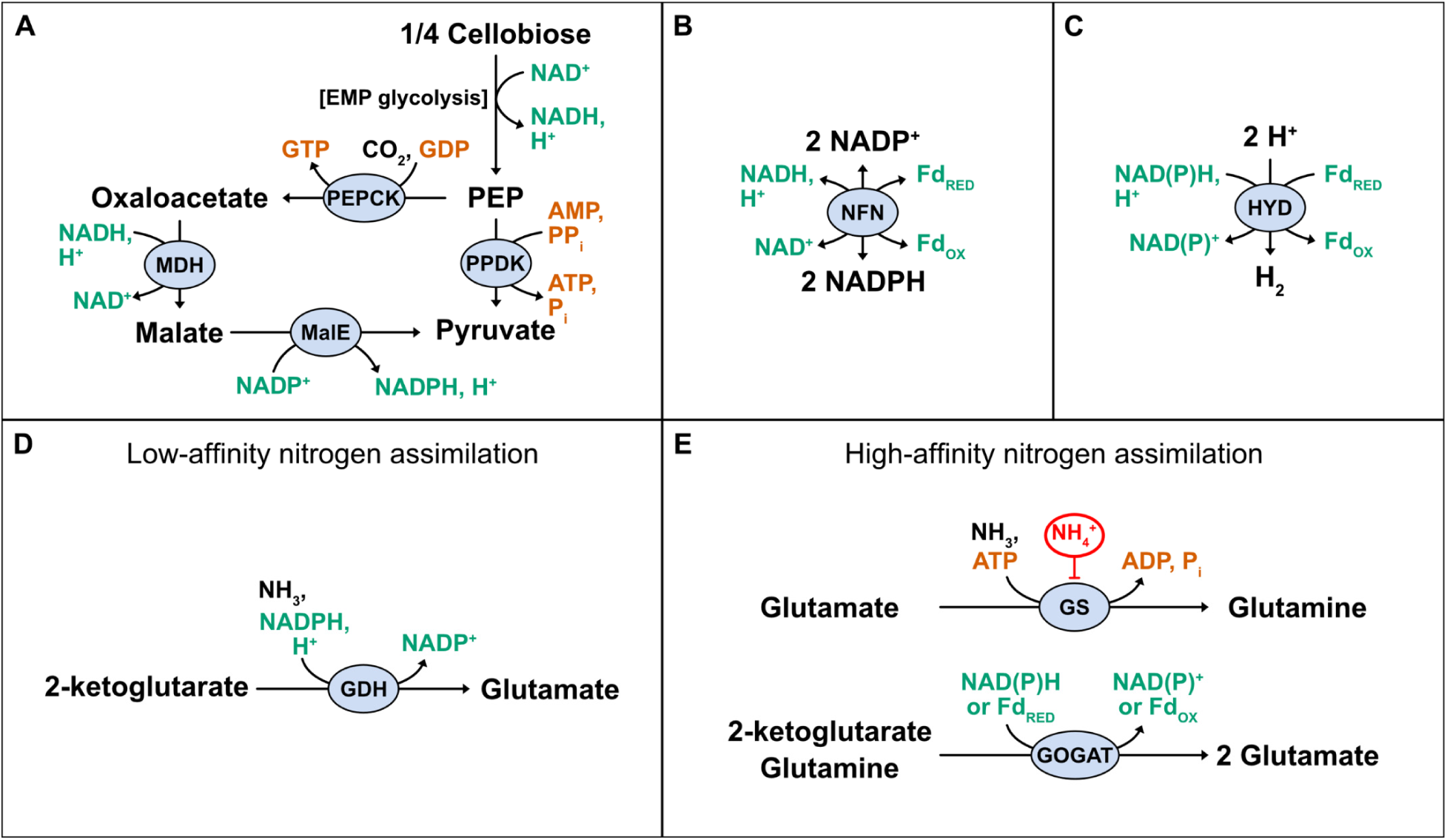
Cofactor usage of *C. thermocellum* in (A) the PEP-to-pyruvate conversion, either via the redox-independent PPDK or the NADPH-producing malate shunt, (B) the NADH-dependent NFN reaction, (C) a summarized HYD reaction, (D) the low-affinity GDH reaction and (E) the high-affinity GS-GOGAT cycle. Abbreviations: EMP, Embden-Meyerhof-Parnas; PEP, phosphoenolpyruvate; PPDK, pyruvate phosphate dikinase; PEPCK, PEP carboxykinase; MDH, malate dehydrogenase; MalE, malic enzyme; NFN, NADH-dependent reduced ferredoxin:NADP^+^ oxidoreductase; HYD, hydrogenase; GDH, glutamate dehydrogenase; GS, glutamine synthetase; GOGAT, glutamate synthase.

**TABLE 1.**
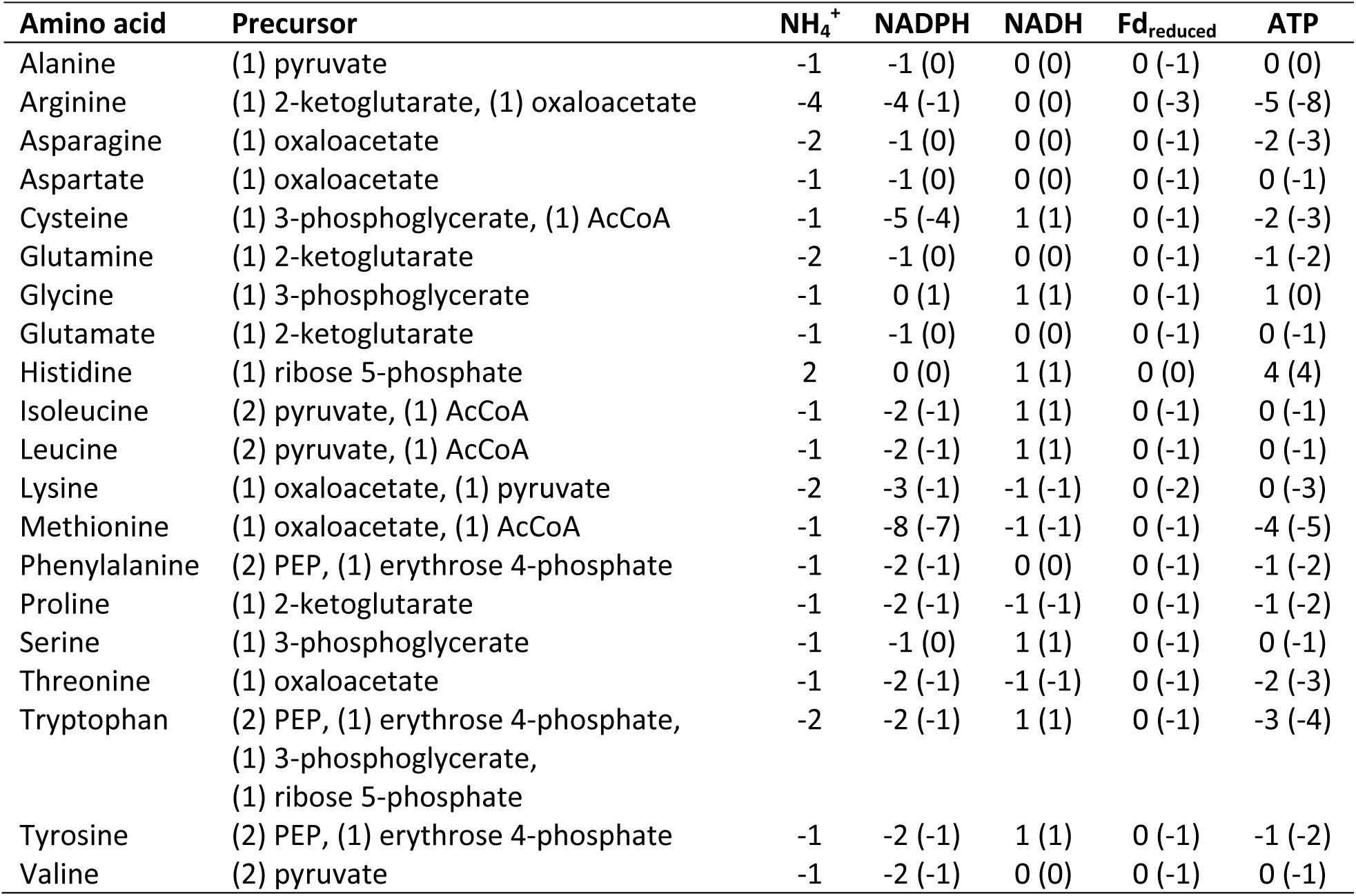
Precursors and stoichiometric coefficients for cofactors used in amino acid synthesis from precursors in C. thermocellum based on the genome-scale metabolic model iCBI655 (11). The numbers in brackets represent cofactor usage if glutamate production occurred via a ferredoxin- and ATP-dependent glutamine synthetase (GS)-glutamate synthase (GOGAT) cycle instead of a NADPH-dependent glutamate dehydrogenase (GDH).

In contrast to many model heterotrophic microorganisms, where the pentose phosphate pathway provides most NADPH, *C. thermocellum* predominantly uses the malate shunt to supply NADPH, by converting phosphoenolpyruvate (PEP) to pyruvate through PEP carboxykinase (PEPCK), NADH-dependent malate dehydrogenase (MDH) and NADP^+^-dependent malic enzyme (encoded by *malE*, Clo1313_1879) (Fig. 1A) (14–17). Alternative to the malate shunt, PEP-to-pyruvate conversion can occur through the alternative non-NADPH-producing (redox-independent) pyruvate phosphate dikinase (encoded by *ppdk*, Clo1313_0949) (Fig. 1A). By changing the flux distribution through the malate shunt with flux through Ppdk, more or less NADPH can be formed. Interestingly, both MalE and Ppdk are activated by ammonium but with different activation constants (K_a_). MalE shows a K_a_ for ammonium of 0.7-0.8 mM (14, 18), whereas Ppdk requires higher ammonium concentration with an estimated K_a_ of 3.8 mM (10). Hence, it has been suggested that at lower intracellular ammonium levels, as is likely under nitrogen-limitation, a flux redistribution between Ppdk and MalE may occur that favors NADPH formation (10). If this flux distribution results in an oversupply of NADPH, increased amino acid production, such as valine, could act as a NADPH sink. *C. thermocellum* also contains an NADH-dependent reduced ferredoxin:NADP^+^ oxidoreductase (encoded by *nfnAB*, Clo1313_1848-1849) (Fig. 1B) that might contribute to formation of NADPH by transferring electrons from NADH and reduced ferredoxin to NADP^+^ or, alternatively, reoxidize excess NADPH by producing reduced-ferredoxin and NADH (12).

In addition to the regulatory role on Ppdk and MalE, (intracellular) ammonium levels commonly regulate the activities of the ammonium-assimilation pathways through glutamate dehydrogenase (GDH) or the higher-affinity glutamine synthetase (GS)-glutamate synthase (GOGAT) (Fig. 1D, E) (19, 20). *C. thermocellum* has genes encoding one NADPH-dependent GDH (Clo1313_1847), four GS (three Type III GS, *glnN*, Clo1313_1357, 2038, 2303; one Type Iα GS, *glnA*, Clo1313_2031), and a GOGAT cluster (*gogat*, Clo1313_2032-2036) (21, 22). Enzyme activities for GDH and GS were confirmed *in vitro* (10, 23). In line with expectations, GS activity in *C. thermocellum* increases at nitrogen-depleted conditions (23). Despite several attempts, GOGAT activity has hitherto not been measured in *C. thermocellum* (23, 24). Failure to measure NADH- or NADPH-dependent activity is in line with the current KEGG genome annotation of a ferredoxin binding-subunit (Clo1313_2035) in the *gogat* operon of *C. thermocellum* (21, 23, 24). This would mean that a switch from the NADPH-dependent GDH (Fig. 1D) to a higher-affinity ferredoxin-linked GS-GOGAT system (Fig. 1E), would decrease the use of NADPH reoxidized in amino acid biosynthesis (Table 1). In the example of valine, the decrease from two to one NADPH (Table 1) would mean that regeneration of the same amount of NADPH would require a doubling of the valine secretion flux, possibly contributing to the observed increased amino acid secretion under nitrogen limitation.

The aim of this study was to investigate the role of NADPH-supplying and -consuming pathways in *C. thermocellum* on amino acid secretion. The contribution of the NADPH-supplying malate shunt was investigated in strains relying fully on either the NADPH-yielding malate shunt (in a Δ*ppdk* strain) or redox-independent conversion of PEP-to-pyruvate (in a Δ*ppdk* Δ*malE::P_eno_-pyk* strain). To investigate another potential NADPH source, a Δ*nfnAB* strain was tested under identical conditions. Finally, the role of the GDH/GS-GOGAT node was investigated by testing a Δ*gogat* strain. All strains were characterized and compared to the reference strain in cellobiose- or ammonium-limited chemostat cultures through determination of substrate, biomass and extracellular metabolite yields at a fixed dilution rate of 0.1 h^-1^. Enzyme activity assays were performed to confirm the targeted gene deletions as well as investigate regulation due to the gene deletions or nutrient limitation.

## RESULTS

### Elimination of the malate shunt decreased amino acid secretion

The contribution of NADPH formation at the PEP-to-pyruvate node to amino acid secretion was investigated by constructing strains that either fully relied on the NADPH-forming malate shunt or on the alternative redox-independent conversion by Pyk. Removal of *ppdk* resulted in a strain fully relying on the NADPH-forming malate shunt (Δ*ppdk*, named AVM003 (25)). Hitherto, construction of a strain relying solely on Ppdk for PEP-to-pyruvate conversion has not been successful (13). To obtain a strain fully relying on redox-independent conversion of PEP-to-pyruvate, the native *malE* gene was swapped with a heterologous pyruvate kinase (*pyk*) from *Thermoanaerobacterium saccharolyticum* (13, 34), in the Δ*ppdk* strain, resulting in strain AVM064 (Δ*ppdk* Δ*malE::P_eno_-pyk*). Using the Δ*ppdk* background for AVM064 simultaneously avoids competition/interference between Ppdk and Pyk and any influence of ammonium activation on Ppdk. Removal of Ppdk and MalE in the respective strains was confirmed by enzyme activity assays and a high Pyk activity was measured in AVM064 (Table 2). The reference strain showed small background Pyk activity that can be attributed to the combined reaction of PEPCK (15) and MDH (Table 2, Fig. 1) (26).

**TABLE 2.** Activities of Ppdk, MalE and heterologously expressed Pyk (T. saccharolyticum) in batch cultures.^a,b^

| Strain | Enzyme activity ($\mu\text{mol mg}_{\text{protein}}^{-1} \text{min}^{-1}$) | | |
| --- | --- | --- | --- |
|  | Ppdk | MalE | Pyk |
| Reference strain (DSM 1313) | $0.28 \pm 0.02$ | $5.8 \pm 0.3$ | $0.08 \pm 0.03$ |
| AVM003 ($\Delta ppdk$) | <0.05 | $12.5 \pm 1.3$ | $0.06 \pm 0.03$ |
| AVM064 ($\Delta ppdk \Delta malE::P_{\text{eno}}-pyk$) | <0.05 | <0.05 | $38.4 \pm 2.2$ |

<sup>a</sup> The detection limit was $0.05 \mu\text{mol mg}_{\text{protein}}^{-1} \text{min}^{-1}$ .
<sup>b</sup> Data are shown as average ± standard deviation for four technical replicates of biological duplicates.

The hypothesis that changes in flux distribution at the PEP-to-pyruvate node, caused by varying intracellular ammonium concentrations, influence amino acid secretion, was investigated using the engineered and wild-type strains in chemostat cultures that were limited in either ammonium (N-source) or cellobiose (C-source). If the previously observed increased amino acid secretion by the wild-type under nitrogen limitation (10) is indeed caused by higher activity and larger flux distribution through the malate shunt compared to Ppdk, resulting in an NADPH oversupply, amino acid secretion should be high and independent of the nutrient limitation in the strain solely relying on the malate shunt, and constitutively low in the strain solely relying on pyruvate kinase. The chemostats were designed with a feed containing 5 g L^-1^ cellobiose and either 0.09 or 0.7 g L^-1^ ammonium to achieve ammonium- or cellobiose-limitation, respectively, based on the rigorous study by Holwerda et al. (10) that evaluated different C/N ratios. The targeted limitation was confirmed by residual cellobiose and ammonium, respectively, being below the detection limit when the respective substrate was limiting and in excess when not limiting (Table 3). As commonly seen for various microorganisms when growth is not limited by the energy source (27–30), ammonium-limited chemostats showed higher cellobiose uptake rates and slightly lower biomass yields on cellobiose (Table 3).

**TABLE 3.** Physiological parameters in chemostats with either cellobiose or ammonium limitation, at a dilution rate of 0.10 h^-1^.^a,b^

|  | Cellobiose-limitation |  |  |  |  | Ammonium-limitation |  |  |  |  |
| --- | --- | --- | --- | --- | --- | --- | --- | --- | --- | --- |
| | Reference strain (LL345) | AVM064 ($\Delta ppdk$ $\Delta malE::P_{eno^-} pyk$) | AVM003 ($\Delta ppdk$) | LL1084 ($\Delta nfnAB$) | AG1715 ($\Delta gogat$) | Reference strain (LL345) | AVM064 ($\Delta ppdk$ $\Delta malE::P_{eno^-} pyk$) | AVM003 ($\Delta ppdk$) | LL1084 ($\Delta nfnAB$) | AG1715 ($\Delta gogat$) |
| **Cellobiose in feed (g L<sup>-1</sup>)** | 4.70 ± 0.03 | 4.98 ± 0.03 | 4.76 ± 0.08 | 4.87 ± 0.02 | 4.80 ± 0.04 | 4.69 ± 0.04 | 4.99 ± 0.00 | 4.63 ± 0.04 | 4.85 ± 0.02 | 4.77 ± 0.06 |
| **Residual cellobiose (g L<sup>-1</sup>)** | <0.01 | <0.01 | <0.01 | <0.01 | <0.01 | 1.83 ± 0.03 | 1.10 ± 0.04 | 1.66 ± 0.11 | 1.57 ± 0.10 | 2.23 ± 0.30 |
| **Ammonium in feed (g L<sup>-1</sup>)** | 0.69 ± 0.00 | 0.71 ± 0.02 | 0.67 ± 0.02 | 0.70 ± 0.01 | 0.70 ± 0.01 | 0.087 ± 0.003 | 0.089 ± 0.001 | 0.087 ± 0.001 | 0.089 ± 0.001 | 0.089 ± 0.001 |
| **Residual ammonium (g L<sup>-1</sup>)** | 0.51 ± 0.01 | 0.53 ± 0.01 | 0.50 ± 0.01 | 0.45 ± 0.01 | 0.45 ± 0.01 | <0.002 | <0.002 | <0.002 | <0.002 | <0.002 |
| **q<sub>Cellobiose</sub> (mmol g<sub>biomass</sub><sup>-1</sup> h<sup>-1</sup>)** | -2.19 ± 0.03 | -2.36 ± 0.08 | -2.07 ± 0.09 | -2.20 ± 0.02 | -2.15 ± 0.02 | -2.49 ± 0.02 | -2.54 ± 0.10 | -2.40 ± 0.10 | -2.45 ± 0.12 | -2.11 ± 0.06 |
| **q<sub>Ammonium</sub> (mmol g<sub>biomass</sub><sup>-1</sup> h<sup>-1</sup>)** | -1.60 ± 0.04 | -1.62 ± 0.07 | -1.56 ± 0.04 | -2.14 ± 0.12 | -2.07 ± 0.06 | -1.43 ± 0.05 | -1.10 ± 0.05 | -1.33 ± 0.10 | -1.26 ± 0.03 | -1.41 ± 0.11 |
| **Biomass yield (g g<sub>cellobiose</sub><sup>-1</sup>)** | 0.13 ± 0.00 | 0.12 ± 0.01 | 0.15 ± 0.01 | 0.13 ± 0.00 | 0.14 ± 0.00 | 0.12 ± 0.00 | 0.11 ± 0.00 | 0.13 ± 0.01 | 0.12 ± 0.01 | 0.14 ± 0.00 |
| **Glycogen content (% g g<sub>biomass</sub><sup>-1</sup>)** | 1.1 ± 0.6 | N.D. | N.D. | N.D. | N.D. | 31.5 ± 0.6 | N.D. | N.D. | N.D. | N.D. |
| **Ethanol yield (mol mol<sub>cellobiose</sub><sup>-1</sup>)** | 0.97 ± 0.06 | 1.55 ± 0.01 | 0.96 ± 0.03 | 0.96 ± 0.02 | 0.84 ± 0.07 | 1.07 ± 0.03 | 1.55 ± 0.05 | 0.84 ± 0.05 | 1.00 ± 0.02 | 0.77 ± 0.03 |
| **Acetate yield (mol mol<sub>cellobiose</sub><sup>-1</sup>)** | 1.38 ± 0.01 | 0.65 ± 0.02 | 1.46 ± 0.04 | 1.38 ± 0.02 | 1.38 ± 0.01 | 0.94 ± 0.01 | 0.61 ± 0.01 | 1.17 ± 0.12 | 1.01 ± 0.01 | 0.84 ± 0.07 |
| **Formate yield (mol mol<sub>cellobiose</sub><sup>-1</sup>)** | 0.11 ± 0.04 | 0.22 ± 0.02 | 0.10 ± 0.01 | 0.31 ± 0.01 | 0.07 ± 0.01 | 0.15 ± 0.01 | 0.32 ± 0.03 | 0.18 ± 0.05 | 0.13 ± 0.03 | 0.30 ± 0.12 |
| **Lactate yield (mol mol<sub>cellobiose</sub><sup>-1</sup>)** | 0.01 ± 0.00 | 0.02 ± 0.00 | 0.02 ± 0.00 | 0.01 ± 0.00 | 0.02 ± 0.00 | 0.07 ± 0.01 | 0.04 ± 0.02 | 0.06 ± 0.02 | 0.06 ± 0.00 | 0.04 ± 0.02 |
| **Pyruvate yield (mol mol<sub>cellobiose</sub><sup>-1</sup>)** | <0.01 | <0.01 | 0.01 ± 0.01 | <0.01 | 0.01 ± 0.00 | 0.09 ± 0.00 | 0.01 ± 0.00 | 0.02 ± 0.00 | 0.03 ± 0.00 | 0.02 ± 0.01 |
| **Malate yield (mol mol<sub>cellobiose</sub><sup>-1</sup>)** | <0.01 | 0.11 ± 0.02 | <0.01 | <0.01 | <0.01 | <0.01 | 0.03 ± 0.00 | <0.01 | <0.01 | <0.01 |
| **Total amino acids yield (mol mol<sub>cellobiose</sub><sup>-1</sup>)<sup>c</sup>** | 0.026 ± 0.001 | 0.031 ± 0.002 | 0.028 ± 0.002 | 0.030 ± 0.001 | 0.022 ± 0.001 | 0.116 ± 0.002 | 0.044 ± 0.006 | 0.107 ± 0.026 | 0.115 ± 0.009 | 0.038 ± 0.010 |
| **CO<sub>2</sub> yield (mol mol<sub>cellobiose</sub><sup>-1</sup>)<sup>d</sup>** | 2.38 ± 0.02 | 2.00 ± 0.04 | 2.48 ± 0.06 | 2.18 ± 0.03 | 2.29 ± 0.05 | 2.07 ± 0.03 | 1.96 ± 0.01 | 2.06 ± 0.08 | 2.11 ± 0.02 | 1.48 ± 0.12 |
| **H<sub>2</sub> yield (mol mol<sub>cellobiose</sub><sup>-1</sup>)** | 4.85 ± 0.06 | N.D. | N.D. | N.D. | 4.73 ± 0.05 | 2.44 ± 0.09 | N.D. | N.D. | N.D. | 4.26 ± 0.46 |
| **Ex. protein yield (mol mol<sub>cellobiose</sub><sup>-1</sup>)<sup>e</sup>** | 0.09 ± 0.00 | 0.06 ± 0.00 | 0.08 ± 0.00 | 0.10 ± 0.00 | 0.08 ± 0.00 | 0.05 ± 0.00 | 0.04 ± 0.00 | 0.05 ± 0.00 | 0.04 ± 0.00 | 0.04 ± 0.01 |
| **Carbon recovery (%)<sup>f</sup>** | 81 ± 1 | 77 ± 0 | 84 ± 2 | 81 ± 1 | 77 ± 1 | 77 ± 1 | 75 ± 1 | 77 ± 3 | 76 ± 1 | 64 ± 1 |
| **Nitrogen recovery (%)<sup>f</sup>** | 71 ± 2 | 66 ± 2 | 70 ± 2 | 54 ± 3 | 53 ± 2 | 87 ± 2 | 95 ± 3 | 96 ± 2 | 97 ± 1 | 72 ± 6 |
| **Degree-of-reduction recovery (%)<sup>f</sup>** | 70 ± 1 | 73 ± 0 | 72 ± 2 | 70 ± 1 | 66 ± 1 | 69 ± 1 | 71 ± 1 | 68 ± 2 | 68 ± 1 | 59 ± 1 |
| **Degree-of-reduction recovery with H<sub>2</sub> (%)<sup>f</sup>** | 90 ± 1 | N.D. | N.D. | N.D. | 86 ± 2 | 79 ± 1 | N.D. | N.D. | N.D. | 76 ± 1 |

<sup>a</sup> Data are shown as average ± standard deviation for three biological replicates.<sup>b</sup> Negative values represent consumption and positive values represent production.<sup>c</sup> Excluding cysteine.<sup>d</sup> Calculated from fermentation product yields and biomass yield (see Materials and Methods).<sup>e</sup> Based on a molecular weight of a generalized protein composition (see Supplemental Table S1 and Materials and Methods).<sup>f</sup> Calculated as the ratio between total element or degree-of-reduction found in products over the total element or degree-of-reduction in consumed substrate (see Supplemental Tables S3-S5).
N.D. not determined.

To create a baseline for subsequent comparisons of the other strains under identical conditions, the reference strain LL345 was grown in cellobiose- or ammonium-limited chemostats (Table 3). The total amino acid yield increased 4.5-fold, from 0.026 ± 0.001 mol mol_cellobiose_^-1^ under cellobiose limitation to 0.116 ± 0.002 mol mol_cellobiose_^-1^ under ammonium limitation (Table 3). Not only did the yields of the pyruvate-derived amino acids alanine (1.5-fold), isoleucine (7.8-fold), leucine (3.1-fold) and valine (20.4-fold) increase, but also the extracellular pyruvate yield (from <0.01 to 0.09 ± 0.00 mmol mol_cellobiose_^-1^) and lactate yield (4.7-fold) increased (Table 3, Fig. 2, Supplemental Table S2). Other amino acid yields that increased were methionine (from undetected to 6.3 ± 0.5 µmol mol_cellobiose_^-1^), phenylalanine (3.9-fold), glutamate (2.1-fold) and tyrosine (2.1-fold) (Fig. 2, Supplemental Table S2). No significant changes were observed for asparagine, aspartate, glutamine, arginine, glycine, histidine, threonine, or tryptophan excretion, whereas the serine yield dropped by 26% (Supplemental Table S2). In absolute numbers, valine, alanine and glutamate each accounted for approximately 15-20% of the secreted amino acids under cellobiose limitation (Fig. 2). Under ammonium limitation, valine was by far the most abundant at 58% of the secreted amino acids (Fig. 2). As a result, secreted amino acids accounted for 4.9% of the carbon and 5.7% of the degree-of-reduction of the consumed cellobiose under ammonium limitation compared to only 0.9% and 0.9%, respectively, under cellobiose limitation (Table S3 and S4). Similarly, 20.3% of the total consumed nitrogen ended up in secreted amino acids in ammonium limitation compared to 3.7% in cellobiose limitation (Table S5). Among the traditional fermentation products, the largest change was observed for the H_2_ yield, which dropped by 50% under ammonium limitation. Additionally, the intracellular glycogen content for the reference strain increased from 1.1 ± 0.6% (w/w) under cellobiose limitation to 31.5 ± 0.6% (w/w) under ammonium limitation. The results of this baseline characterization of the reference strain in cellobiose-/ammonium-limited chemostats are consistent with the previous results of Holwerda et al. (10). Under identical conditions, the strain fully relying on Pyk for conversion of PEP-to-pyruvate (AVM064; Δ*ppdk* Δ*malE::P_eno_-pyk*), thereby lacking the NADPH provided by the malate shunt, only showed a 1.4-fold increase (*P* < 0.05) in the total amino acid yield from cellobiose limitation to ammonium limitation, compared to 4.5-fold observed for the reference strain (Table 3). Although valine secretion increased 10-fold (*P* < 0.05) from cellobiose to ammonium limitation, the valine yield under ammonium limitation was still 81% lower than the reference strain (Fig. 2, Supplemental Table S2). Increases in glutamate (1.6-fold), isoleucine (5.6-fold), leucine (1.6-fold) and tyrosine (1.4-fold) yields (*P* < 0.05) under ammonium limitation were also lower than that for the reference strain by 28%, 63%, 8%, and 40%, respectively (*P* < 0.05) (Fig. No significant differences between the two limitations for AVM064 were observed for alanine and lactate (Fig. 2), whilst pyruvate increased from <0.01 to 0.01 ± 0.00 mol mol_cellobiose_^-1^ (*P* < 0.05), which was 87% smaller than the reference strain in ammonium limitation (Table 3). Overall, rerouting the PEP-to-pyruvate conversion through Pyk rather than the malate shunt decreased the total carbon, degree-of-reduction and nitrogen ending up in the secreted amino acids under nitrogen limitation to 1.9%, 2.1%, and 10.8%, respectively, compared to 4.9%, 5.7% and 20.3% for the reference strain under the same conditions (Supplemental Tables S3-S5). The malate shunt in AVM064 was interrupted through removal of malic enzyme, whilst maintaining malate dehydrogenase, resulting in secretion of malate with a yield of 0.11 ± 0.02 mol mol_cellobiose_^-1^ in cellobiose limitation and 0.03 ± 0.00 mol mol_cellobiose_^-1^ in ammonium limitation (Table 3). In line with a switch from a malate shunt, which transfers electrons from NADH to NADP^+^, to the redox-independent PEP-to-pyruvate conversion in the Pyk-dependent strain, the ethanol yield increased by 46-60% to 1.55 mol mol_cellobiose_^-1^ in either nutrient limitation compared to the reference strain (Table 3).

**FIG. 2.**
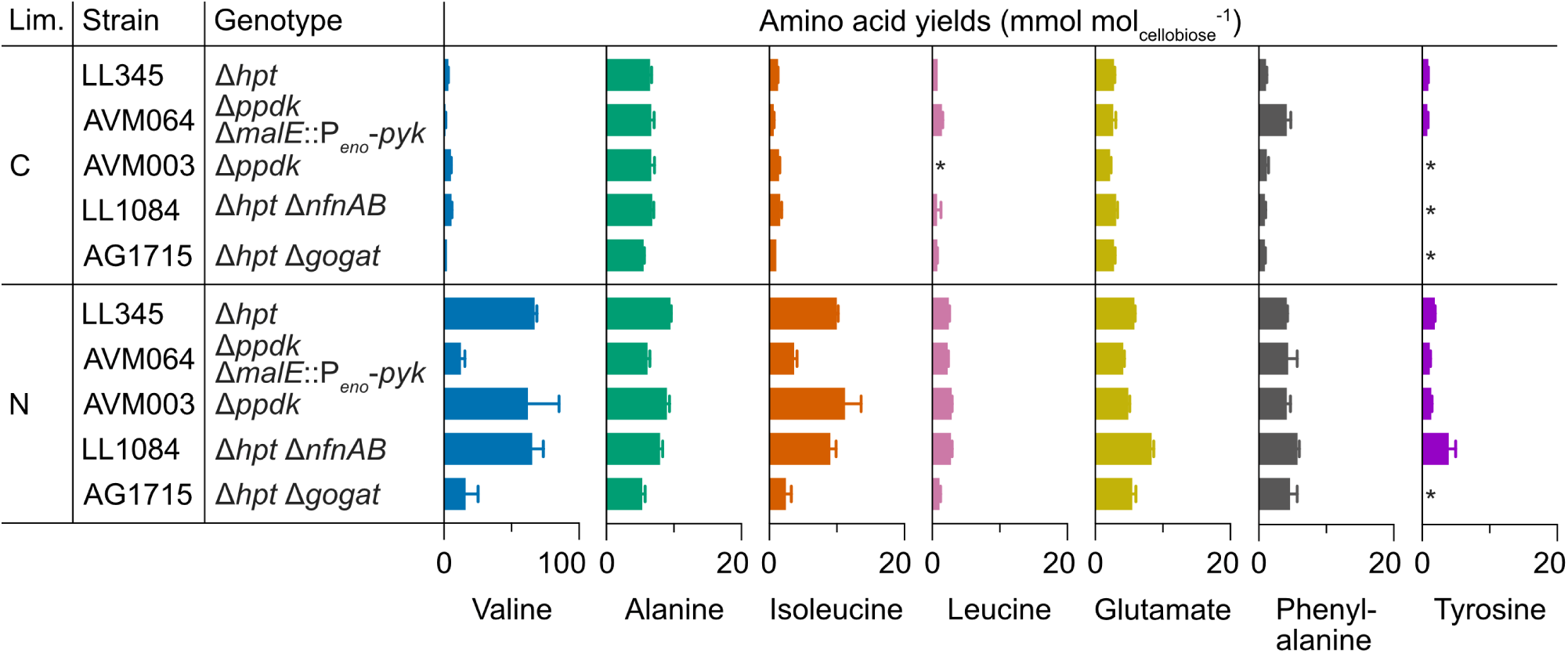
Amino acid yields in chemostats with either cellobiose (C) or ammonium (N) as the sole limiting nutrient, at a dilution rate of 0.10 h^-1^. The remaining amino acids can be found in Supplemental Table S6 and were either below detection limit (*, <0.5 mmol mol_cellobiose_^-1^) or only showed small changes. Error bars represent standard deviation for three biological replicates.

The physiological impact of deleting *ppdk*, thereby routing the entire PEP-to-pyruvate conversion through the NADPH-yielding malate shunt, did not result in any drastic changes in product yields compared to the reference strain (Table 3). In the ammonium-limited chemostat cultures, the impact of *ppdk* deletion was slightly higher, with a 21% decrease in the ethanol yield from 1.07 ± 0.03 mol mol_cellobiose_^-1^ in the reference strain to 0.84 ± 0.05 mol mol_cellobiose_^-1^ in the Δ*ppdk* strain, and a corresponding increase in the acetate yield (Table 3). Amino acid yields in both limitations were similar to the reference strain (Fig. 2, Supplemental Table S2).

The lack of a physiological impact of the *ppdk* deletion in the cellobiose-limited chemostats (Table 3) suggests that the flux through Ppdk in wild-type under this condition was likely insignificant. Even though NADPH supply through the malate shunt is clearly required for the surplus amino acid secretion under ammonium-limited conditions, as seen from the strongly decreased total amino acid yield observed for AVM064 (Δ*ppdk* Δ*malE*::P*_eno_*-*pyk*) (Table 3), this suggests a more complex underlying mechanism than only ammonium-dependent redistribution of the PEP-to-pyruvate conversion fluxes between Ppdk and MalE.

### No impact of *nfnAB* deletion on amino acid secretion

To investigate the role of the NADH-dependent reduced ferredoxin:NADP^+^ oxidoreductase (NfnAB) (Fig. 1B) in amino acid secretion, a Δ*nfnAB* strain (LL1084) (12) was characterized in cellobiose- or ammonium-limited chemostats. A comparison of this strain with the reference strain in cellobiose-limited chemostats showed the same biomass yield and cellobiose uptake rate in both conditions (Table 3). Under cellobiose limitation, there was a 2.7-fold increase (*P* < 0.05) in formate yield compared to the reference strain, at identical acetate and ethanol yields (Table 3). This suggests a small shift from PFOR, which reduces ferredoxin, to pyruvate formate-lyase (PFL) in the Δ*nfnAB* strain. Interestingly, under ammonium limitation, the formate yield was the same as the reference strain, suggesting negligible flux through NfnAB in the reference strain under this condition or full functional complementation by the other NADPH supplying routes. In line with this, deletion of *nfnAB* also did not result in a meaningful change in total amino acid formation (Fig. 2). Similar to cellobiose-limited chemostats, ammonium limitation of the Δ*nfnAB* strain showed the same biomass yield and cellobiose uptake rate as the reference strain (Table Only a minor decrease of the pyruvate yield from 0.09 ± 0.00 mol mol_cellobiose_^-1^ in the reference strain to 0.03 ± 0.00 mol mol_cellobiose_^-1^ in the Δ*nfnAB* strain, and an accompanying increase of the acetate yield by the same magnitude (*P* < 0.05), were observed.

### Deletion of *gogat* decreased amino acid secretion under nitrogen limitation

To investigate if GOGAT upregulation, and the predicted accompanying shift from NADPH-dependent to partially ferredoxin-linked ammonium assimilation, contribute to amino acid secretion under nitrogen limitation, a Δ*gogat* strain (AG1715) was constructed and characterized in cellobiose- or ammonium-limited chemostats. Under cellobiose limitation, where both strains showed high NADPH-linked GDH activities (25.0 ± 2.5 and 32.3 ± 1.3 µmol mg_protein_^-1^ min^-1^) (Table 4) and GDH is expected to carry the majority of the ammonium-fixation flux, the Δ*gogat* strain and the reference strain showed the same yields and rates for canonical fermentation products (Table 3). Under these conditions, a small decrease in total amino acid secretion of 15% (*P* < 0.05) was observed in the Δ*gogat* strain compared to the reference strain (Table 3).

**TABLE 4.** Glutamate dehydrogenase activities in LL345 and AG1715 from steady-state cultures grown in cellobiose- or ammonium-limited chemostats, expressed in µmol mg_protein_^-1^ min^-1^.^a^

|  | Cellobiose limitation |  | Ammonium limitation |  |
| --- | --- | --- | --- | --- |
| | Reference strain (LL345) | AG1715 ($\Delta\text{gogat}$) | Reference strain (LL345) | AG1715 ($\Delta\text{gogat}$) |
| **GDH** |  |  |  |  |
| - NADPH | 25.0 ± 2.5 | 32.3 ± 1.3 | 0.39 ± 0.04 | 62.3 ± 10.3 |
| - NADH | 0.16 ± 0.01 | 0.38 ± 0.03 | <0.05 | 0.36 ± 0.02 |

<sup>a</sup> Data are shown as average ± standard deviation for four technical replicates of biological duplicates.

In ammonium-limited chemostats, the GDH activity of the reference strain dramatically decreased 64-fold to 0.39 ± 0.04 µmol mg_protein_^-1^ min^-1^ (Table 4), which is in line with a text-book switch to the higher affinity GS-GOGAT system (activity not measured). With GDH being the sole ammonium-assimilating option in the Δ*gogat* strain, GDH activity was upregulated 160-fold to 62.3 ± 10.3 µmol mg_protein_^-1^ min^-1^ compared to the reference strain under ammonium limitation (Table 4). Interestingly, comparing the Δ*gogat* strain between growth conditions also showed a 2-fold upregulation (*P* < 0.05) of GDH activity under ammonium limitation. This upregulation was likely needed to compensate for the (predicted) lower intracellular ammonium concentration and to maintain the same ammonium-assimilation flux needed at the identical growth rates of 0.10 h^-1^.

In ammonium-limited chemostats, the total amino acid secretion decreased by 67%, from 0.116 ± 0.002 mol mol_cellobiose_^-1^ in the reference strain to 0.038 ± 0.010 mol mol_cellobiose_^-1^ in the Δ*gogat* strain (Table 3). Valine decreased the most and had a yield of 16.0 ± 9.4 mmol mol_cellobiose_^-1^ in ammonium limitation, which was 76% lower than the reference strain (*P* < 0.05) (Fig. 2). Yields of other pyruvate-derived amino acids also significantly decreased compared to the reference strain, with 44% lower alanine, 75% lower isoleucine and 56% lower leucine in ammonium limitation (*P* < 0.05) (Fig. 2). All remaining amino acid yields, except that of phenylalanine, were significantly lower in the Δ*gogat* strain (*P* < 0.05) (Fig. 2, Supplemental Table S2). The relative abundance of valine dropped from 58% of total secreted amino acids in the reference strain to 42% in the Δ*gogat* strain (Supplemental Table S2). Phenylalanine, glutamate and alanine followed and corresponded to roughly 12%, 15%, and 14%, respectively. Overall, the deletion of *gogat* decreased the amount of carbon, degree-of-reduction, and nitrogen consumed diverted to the amino acid byproducts under ammonium limitation from 4.9%, 5.7% and 20.3% in the reference strain to 1.6%, 1.8%, and 5.9% in the Δ*gogat* strain (Supplemental Tables S3-S5).

Surprisingly, deletion of *gogat* resulted in a large increase of the H_2_ yield in ammonium limitation. The reference strain had a H_2_ yield of 2.44 ± 0.09 mol mol_cellobiose_^-1^, whereas the Δ*gogat* strain showed a H_2_ yield of 4.26 ± 0.46 mol mol_cellobiose_^-1^ (Table 3), which might reflect an increased need for ferredoxin reoxidation through the hydrogenases. Concomitantly, the ethanol yield decreased with 0.3 mol mol_cellobiose_^-1^ (27%) (*P* < 0.05), whereas the formate yield doubled to 0.30 ± 0.12 mol mol_cellobiose_^-1^ (*P* < 0.05) (Table 3). These changes in redox metabolism would be in line with the increased stoichiometric need for NADPH in the ammonium assimilation if the NADPH that is formed through the malate shunt goes at the expense of NADH. The biomass yield increased 19% compared to the reference strain, which at least directionally can be explained by the ATP savings made in biosynthesis when shifting from ammonium assimilation via the ATP-consuming GOGAT-GS cycle to the ATP-neutral GDH reaction.

## DISCUSSION

Using targeted gene deletions and cellobiose-/ammonium-limited chemostats as diagnostic tools, this study found several indications that amino acid secretion is driven by a cellular need to balance NADPH. This was illustrated by decreased amino acid yields upon eliminating the NADPH-supplying malate shunt and separately by likely changing the cofactor specificity of ammonium assimilation. Although a hypothesis of differential regulation of the malate shunt and Ppdk by the ammonium concentration was falsified, the mechanisms underlying amino acid secretion clearly involve oversupply of NADPH that, due to transhydrogenation in the malate shunt, comes at the expense of NADH supply. The lower NADH supply and an apparent minimal contribution of NfnAB for NADPH-to-NADH conversion might limit conversion of pyruvate to lactate and ethanol and instead favor production of NADPH-linked, pyruvate-derived amino acids. Using amino acid formation to regenerate NADP^+^ has been observed in other species as well. This mechanism has been proposed in the fungus *Aspergillus nidulans* (31) and in the alanine-producing archaeon *Pyrococcus furiosus* (32). Anaerobic rumen bacteria also commonly produce amino acids that regenerate NADP^+^ at low ATP cost (33). Amino acid secretion is found among other Clostridia species as well (28, 34), which generally have 5 to 20-fold higher intracellular amino acid levels than Gram-negative bacteria (35).

In addition to the NADPH originating from the malate shunt, specifically under ammonium limitation, a shift from GDH to GOGAT likely decreases the NADPH reoxidized per amino acid and thereby increases amino acid secretion in wild-type *C. thermocellum* under these laboratory conditions. Underlying that hypothesis is the proposed use of ferredoxin as the redox cofactor of GOGAT. This currently unconfirmed annotation (21) is supported by the observed changes in the redox metabolism of the reference strain and Δ*gogat* strain under ammonium limitation (Table 3, Fig. 2). The drop in H_2_ yield in ammonium limitation by the reference strain (Table 3) is in line with a switch from (mainly) NADPH-dependent GDH activity in cellobiose limitation to (mainly) ferredoxin-linked GS-GOGAT activity in ammonium assimilation, which decreases the availability of reduced ferredoxin for H_2_ formation. In addition to this switch, a 17% decrease in production of reduced ferredoxin by pyruvate ferredoxin oxidoreductase (PFOR), (Table 3; calculated as *Y*_acetate/cellobiose_ + *Y*_ethanol/cellobiose_ − *Y*_formate/cellobiose_) is likely also contributing to the drop in H_2_ yield. The Δ*gogat* strain would likely require other mechanisms to reoxidize ferredoxin, which in *C. thermocellum* can (in theory) occur through: *i*) H_2_ production via a proton-pumping energy-converting hydrogenase (ECH) or a bifurcating NAD(P)H-HYD, *ii*) proton-pumping RNF, which increases the NADH pool, *iii*) NfnAB, by consuming NADH and producing NADPH, or *iv*) a cycle with reverse PFOR and forward PFL, which assimilates CO_2_ and generates formate (12). The decreased ethanol yield (suggesting lower NADH availability), increased formate yield, decreased amino acid yields and increased H_2_ yield of the Δ*gogat* strain under ammonium limitation suggest that the combination of HYDs and a 40% lower net flux through PFOR (calculated as *q*_acetate_ + *q*_ethanol_ − *q*_formate_, Supplemental Table S6) replace GOGAT as a ferredoxin sink (Table 3, Fig. 2).

Current stoichiometric metabolic models of *C. thermocellum* contain ‘free’ shuffling of electrons between NAD(P) and ferredoxin *via* the transhydrogenases, NFN and RNF, ultimately resulting in H_2_ instead of amino acids as an electron sink (11, 36). However, metabolic flux and thermodynamic analysis suggest a narrow window for thermodynamically favorable conversion *via* the (trans)hydrogenases (37). The lack of a significant effect on the fermentation product profile of Δ*nfnAB* in chemostat cultures in the present study, and in batch cultures in the study of Lo et al. (12), suggests a minimal contribution or full complementarity of NFN activity under the tested conditions in these strain backgrounds. In laboratory environments, H_2_ production via HYDs is likely further thermodynamically limited by H_2_ supersaturation due to poor liquid-to-gas transfer (38). Physiological predictions and design of metabolic engineering strategies would benefit from including these thermodynamic limitations into the models.

The amino acid secretion observed under these laboratory conditions might be less common in natural environments of cellulolytic Clostridia, since these niches are not only deficient in bioavailable nitrogen, but commonly also contain H_2_-consuming methanogens (39, 40). In the presence of these methanogens, hydrogen partial pressures are maintained low, which makes H_2_ production as an electron sink more favorable (39, 40). Indeed, co-cultures with H_2_-consuming methanogens result in higher acetate-to-ethanol ratio (40), whereas increasing the (dissolved) H_2_ partial pressure results in higher ethanol-to-acetate ratio, decreased H_2_ yield, and reverse NAD(P)H-HYD activity (41–43). Sparging laboratory cultures with N_2_, which is believed to make H_2_ production more favorable, shifted the fermentation profile with increases in ethanol and acetate titers and a decrease in valine titer (37). On top of thermodynamic limitations, HYDs are complex organometallic proteins, requiring specific systems for post-translational assembly (13, 44). Hence, their synthesis might consume more ATP than proteins involved in the valine and alanine pathways. Such kinetic limitations were observed with *Clostridium cellulolyticum*, where chemostat cultures at varying dilution rate showed that the regeneration of cofactors through hydrogen gas formation was too slow at high carbon fluxes, resulting in more lactate and polysaccharides (27, 45). Importantly, diversion of electrons to either hydrogen and/or methane in mixed cultures goes at the expense of the ethanol yield on substrate, necessitating a choice between lower theoretical ethanol yields or continued engineering of *C. thermocellum* for improved product yield.

Aiming to get closer to the maximum theoretical ethanol yield would require further metabolic engineering to decrease the 10% of consumed carbon that is currently diverted to amino acids in the *C. thermocellum* strain with the highest yield (6). One strategy to reduce amino acid secretion would be to simplify the redox metabolism by reducing the diversity in redox cofactors. Similar to *Saccharomyces cerevisiae*, relying solely on NADH for glycolysis and ethanol production, this would likely reduce the NADPH-driven amino acid secretion observed in *C. thermocellum*. As a first step, this study has shown that deleting the NADPH-supplying MalE (Δ*ppdk* Δ*malE*::P_eno_-*pyk*) increased the ethanol and malate yields, whereas amino acid yields in chemostat cultures dropped, reflecting a higher NADH and lower NADPH availability. This is in line with the study by Olson et al. (26), where a similarly constructed strain (Δ*ppdk* Δ*malE*-*mdh* Δ*ldh*::P*_eno_*-*pyk*) in batch cultures showed a 3-fold increased ethanol production, whilst valine production dropped 3-fold. However, engineering of the steps required to convert pyruvate to ethanol in a scenario with simplified redox metabolism, through either i) PFOR, AdhE, and RNF, or ii) a pyruvate decarboxylase (PDC: pyruvate → acetaldehyde + CO_2_) and NADH-dependent alcohol dehydrogenase, has had limited success in achieving sufficiently high ethanol yield and titer and require more research (46–48). Alternatively, the ethanol production pathway could be changed to reoxidize the NADPH that is otherwise diverted to amino acid secretion. For instance, laboratory evolution of a Δ*pta* Δ*ldh* strain resulted in a point mutation in the bifunctional alcohol/aldehyde dehydrogenase (*adhE*) that allowed the use of NADPH on top of NADH as cofactor, thereby producing more ethanol and less amino acids (5, 7, 10). Combining above interventions with a deletion in glutamate synthase and/or *gogat* might further reduce amino acid secretion. In line with that, Rydzak et al. (22) showed that deletion of the GS gene *glnA*, increased the ethanol yield and reduced amino acid secretion significantly. Overall, these findings can guide engineering to decrease amino acid secretion and improve the ethanol yield of *C. thermocellum* for sustainable consolidated bioprocessing of lignocellulosic biomass.

## MATERIALS AND METHODS

### Strains and maintenance

The *C. thermocellum* DSM 1313 wild-type (WT) strain was purchased from the DSMZ microorganism collection (www.dsmz.de). All strains used or constructed in this study are listed in Table 5. Freezer stocks were prepared by growing strains in the complex medium CTFUD (described by Olson & Lynd (49)) at 55°C to an OD of 0.6-2.0. After addition of glycerol to 25% (vol/vol), 1 mL aliquots were stocked in cryo-vials (VWR, Stockholm, Sweden) at −80°C. Strain construction and stocking was performed in an anaerobic vinyl chamber from Coy Laboratory Products (TG Instruments, Helsingborg, Sweden) with an atmosphere of 5% H_2_, 10% CO_2_ and 85% N_2_ (Strandmöllen AB, Ljungby, Sweden).

**TABLE 5.** Strains used in this study.

| Strain name | Parent strain | Genotype | Reference |
| --- | --- | --- | --- |
| Wild-type (WT) | - | DSM 1313 | DSMZ<sup>a</sup> |
| LL345 or M1354 | WT | DSM 1313 $\Delta\text{hpt}$ | (60) |
| AVM003 | WT | DSM 1313 $\Delta\text{ppdk}$ | (25) |
| AVM064 | AVM003 | DSM 1313 $\Delta\text{ppdk}$ $\Delta\text{malE}::\text{P}_{\text{eno-pyk}}^b$ | This study |
| LL1084 | LL345 | DSM 1313 $\Delta\text{hpt}$ $\Delta\text{nfnAB}$ | (12) |
| AG1715 | LL345 | DSM 1313 $\Delta\text{hpt}$ $\Delta\text{gogat}$ (Clo1313_2032-2036) | This study |

<sup>a</sup>DSMZ, German Collection of Microorganisms and Cell Cultures GmbH, Braunschweig, Germany (www.dsmz.de).
<sup>b</sup>The *pyk* gene (*tsac\_1363*) is from the *Thermoanaerobacterium saccharolyticum* JW/SL-YS 485 genome.

### Plasmid and strain construction

Plasmids and primers used in this study are listed in Tables 5 and 6. Plasmid construction and propagation were performed in *Escherichia coli* Top10 (*dam*^+^ *dcm*^+^) (Invitrogen, Carlsbad, CA) and BL21 (*dam*^+^ *dcm*^-^) (New England Biolabs, Ipswitch, MA) aerobically in LB medium supplemented with 12-25 µg mL^-1^ chloramphenicol. Purification of plasmid DNA, genomic DNA, and PCR products were performed with commercially available kits from GeneJET (Thermo Fisher Scientific) and QIAprep (QIAGEN, Germantown, MD). Primers were purchased from Invitrogen (Thermo Fischer Scientific). The plasmids were designed for markerless gene deletion according to Olson & Lynd (49) with pDGO145 as a backbone (50). Phusion High-Fidelity DNA polymerase (Thermo Scientific) was used to amplify fragments from *C. thermocellum* genomic DNA, corresponding to 500-1000 bp upstream (5’-flank), downstream (3’-flank), and internal (int) regions of coding sequences of interest (*malE*, Clo1313_1879; *gogat*, Clo1313_2032-2036) as well as the promoter for the enolase gene (178 bp upstream of Clo1313_2090 on the reverse strand) (51), with the primers listed in Table 7. The *pyk* gene (Tsac_1363) was amplified from genomic DNA of *T. saccharolyticum* JW/SL-YS 485 (DSM 8691) using primers 0553 and 0564. The fragments were assembled into the backbone according to the Gibson protocol (52), generating plasmids pJY19 (simultaneous deletion of *malE* and insertion of P*_eno_*-*pyk*) and pNJ22::GOGAT_del (Δ*gogat*). For simultaneous deletion and insertion, the P*_eno_*-*pyk* gene was placed between the 5’-flank and 3’-flank (53). Plasmids were verified by diagnostic PCR and sequencing. Finally, propagation of plasmids in *E. coli* BL21 (*dam*^+^ *dcm*^-^) ensured correct methylation before transformation into *C. thermocellum* (54). Transformation and selection for markerless gene editing in *C. thermocellum* were performed according to Olson & Lynd in CTFUD medium (49). *malE* was deleted while simultaneously inserting P*_eno_*-*pyk* at the *malE* locus in strain AVM003 (Δ*ppdk*), generating AVM064 (Δ*ppdk* Δ*malE::*P*_eno_*-*pyk*). The *gogat* gene cluster was deleted in LL345 (Δ*hpt*) using pNJ22::GOGAT_del, generating strain LL1715 (Δ*hpt* Δ*gogat*). Sequencing of target loci and the 16S rRNA locus using primers listed in Table 7 was performed to confirm correct gene edits and culture purity.

**TABLE 6.** Plasmids used in this study.

| Plasmid name | Description | Genbank Accession no. | Source or reference |
| --- | --- | --- | --- |
| pDGO145 | Plasmid backbone for deletion/insertion | KY852359 | (50) |
| pJY19 | Simultaneous deletion of *malE* (Clo1313_1879) and insertion of $\text{P}_{\text{eno-pyk}}$ | OP441060 | This study |
| pNJ22::GOGAT_del | Deletion of *gogat* (Clo1313_2032-2036) | OP441061 | This study |

**TABLE 7.**
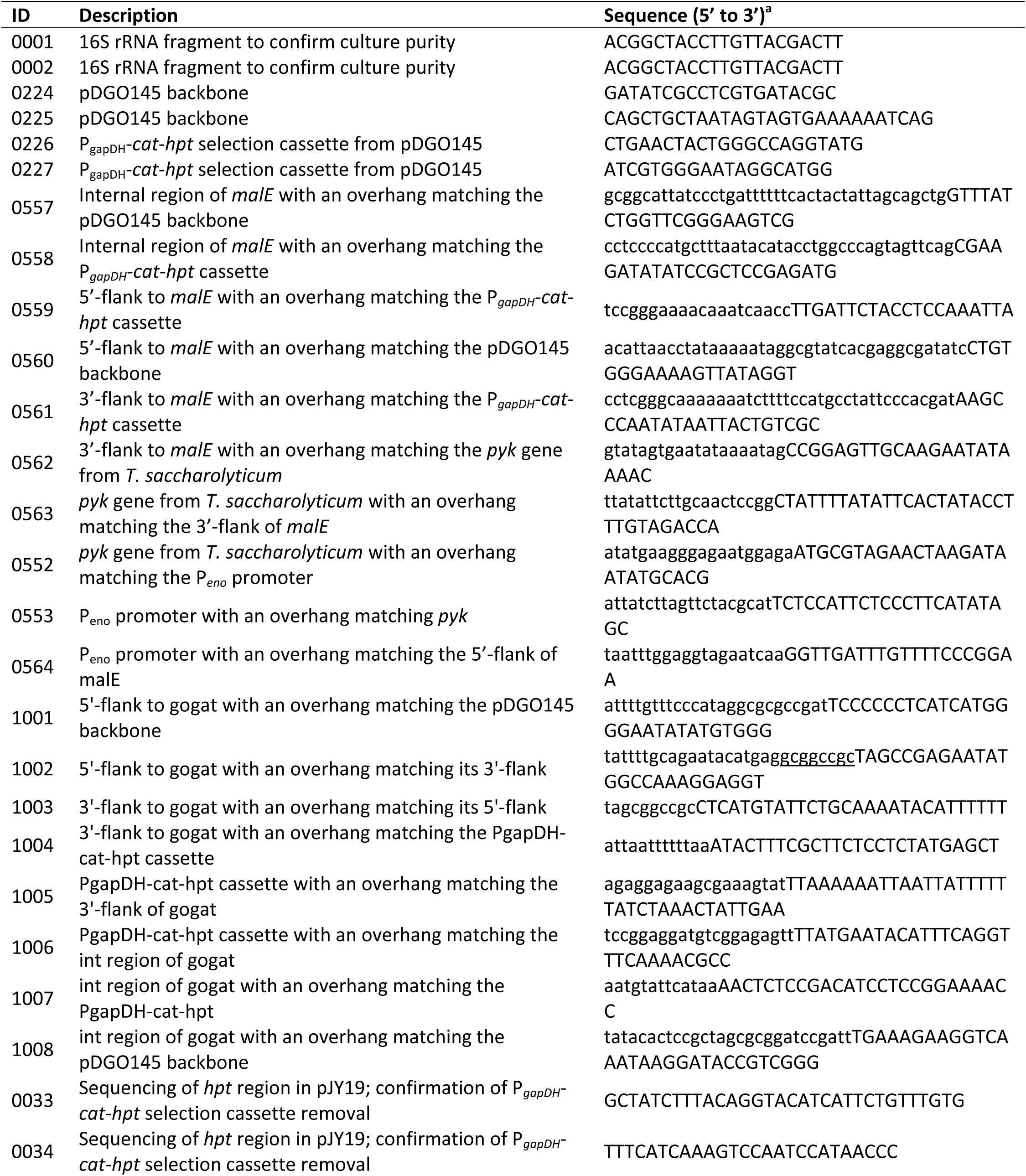
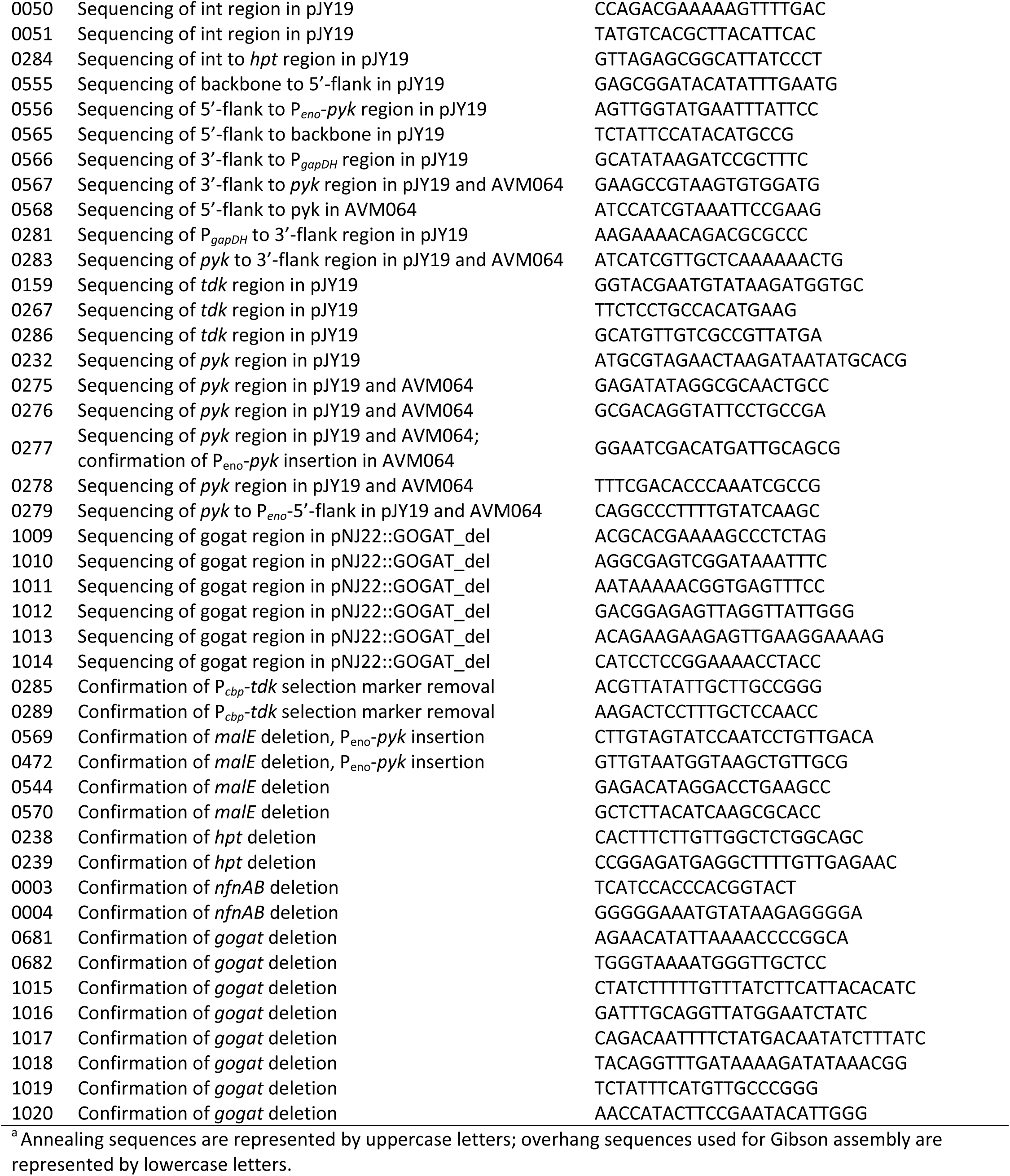
Primers used in this study.

### Media and culture conditions

Serum bottle cultivations were carried out in 125-mL Wheaton serum bottles (DWK Life Sciences, Millville, NJ, USA), closed with blue butyl rubber stoppers (#CLS-4209-14, Chemglass Life Sciences, NJ, US) and contained 50 mL of a defined low-carbon (LC) medium (53, 55). Per liter, the medium contained 5 g D-(+)-cellobiose, 0.5 g urea, 5 g MOPS, 2 g KH_2_PO_4_, 3 g K_2_HPO_4_, 0.1 g Na_2_SO_4_, 0.2 g MgCl_2_·6H_2_O, 0.05 g CaCl_2_·2H_2_O, 0.0035 g FeSO_4_·7H_2_O, 0.025 g FeCl_2_·4H_2_O, 0.1 g L-cysteine-HCl-monohydrate, vitamins (0.02 g pyrodoxamine dihydrochloride, 0.004 g p-aminobenzoic acid, 0.002 g D-biotin, and 0.002 g vitamin B12), and trace elements (1.25 mg MnCl_2_·4H_2_O, 0.5 mg ZnCl_2_, 0.125 mg CoCl_2_·6H_2_O, 0.125 mg NiCl_2_·6H_2_O, 0.125 mg CuSO_4_·5H_2_O, 0.125 mg H_3_BO_3_, and 0.125 mg Na_2_MoO_4_·2H_2_O). The suppliers of chemicals are listed in Yayo et al. (53). The final medium for closed-batch serum bottles was prepared from sterile anaerobic stock solutions as described by Kuil et al. (25) and purged with 20% CO_2_ and 80% N_2_ (Strandmöllen AB) prior to inoculation by alternating vacuum and gas for five cycles.

For chemostats, the LC medium described above was used except with 1 g L^-1^ cysteine, no MOPS and five times higher trace element concentrations, as described by Yayo et al. (53), and the nitrogen source was changed from urea to ammonium to avoid cyclic CO_2_ production observed in preliminary carbon-limited chemostats with urea (at a dilution rate of 0.1 h^-1^). With urea in the feed, a spike in the off-gas CO_2_ concentration every 30 h was found to be coupled to alternating breakdown and accumulation of urea and ammonium (data not shown). Instead, the chemostat feed contained 0.25 g L^-1^ or 2 g L^-1^ ammonium chloride (equal to 0.07 or 0.9 g L^-1^ ammonium) (Sigma) for ammonium- or cellobiose-limited cultivations, respectively (10). The final medium (10 L) was prepared in 10-L Duran flasks (Saveen & Werner AB, Malmö, Sweden) from stock solutions as described by Yayo et al. (53). The feed vessels were covered in aluminum foil and continuously stirred and sparged with filtered N_2_ gas (99.999%) (Nippon Gases, Köping, SE or Air Liquide Gas AB, Malmö, Sweden) at room temperature.

Inocula for serum bottle cultivations and chemostats were prepared by two transfers from the freezer stock in serum bottles. First, an overnight culture was inoculated into 50 mL of fresh medium. At OD 0.5-1.0, this pre-culture was transferred to the main cultivation (serum bottle or bioreactor) with an inoculation volume of 5%. All serum bottles were incubated in a Jeio Tech ISS-4075R incubator (Milmedtek AB, Karlskrona, Sweden) at 55°C and 180 rpm.

### Chemostats

Chemostats were performed in a multi-parallel stainless-steel bioreactor system (GRETA from Belach AB, Stockholm, Sweden), with up to six unpressurized bioreactors running simultaneously at 55°C with 400 rpm agitation. The working volume was maintained at 0.8 L by feedback regulation on the effluent pump via a level sensor. The head space of 0.45 L was continuously purged with filtered N_2_ gas (N5.0, Nippon Gases or Air Liquide Gas) at 0.2 L min^-1^ and the outgas passed through a condenser. In order to minimize O_2_ permeability, Viton O-rings, Viton septa and black Tygon tubing (A-60-G) were used (Sigma-Aldrich). pH was measured with a SteamLine pH electrode (SL80-225pH from VWR) and controlled at 7.0 by addition of filter-sterilized 4.0 M KOH (0.2 µm PES filters) (VWR). The base vessel and 10-L feed vessels were connected via sterile 0.6-mm needles through septa on the head-plate (one feed vessel per reactor). A feed rate of 0.08 L per min, corresponding to a dilution rate of 0.10 h^-1^, was set and resulted in minimal intervals between medium droplets, thereby avoiding feast-famine cycles. The flow rate was checked daily with *in line* glass serological pipettes (25-mL) connected to the feed tube via a T-connection. Effluent was collected from a steam-sterilized sample valve in the bottom of the bioreactor into sterilized plastic Nalgene bottles (VWR) equipped with a gas exhaust filter. Septa were cleaned with 70% isopropanol before and after inoculation, sampling, and addition of feed and base tubes.

A batch cultivation preceded the chemostat phase in the bioreactors and was monitored by base titration or offline optical density (OD) measurements (see below). The feed pump, effluent pump and level sensor were started at the end of the batch or shortly thereafter.

Steady state was defined as a change of less than 5% in biomass concentration over 3 residence times and after at least 4 residence times from feed start. Pre-steady state samples (35 mL) were taken in order to measure OD and cell dry weight (CDW, see below) and thereby determine if steady state had been reached. After this, steady state samples (described below) were taken (9-12 residence times after feed start).

To investigate chemostat homogeneity, OD and CDW of both the reactor and the effluent were compared. No significant differences (>5%) were observed for any of the characterized strains or conditions.

The culture purity of steady-state cultures was assessed by microscopy and sequencing of the 16S rRNA locus, using primers 0001 and 0002 (Table 7). Cross-contamination was excluded by targeted PCR amplification of the strain-specific genotypes.

### H_2_ analysis

The headspace H_2_ mole fraction was measured by offline mass spectrometry. Samples were collected for each bioreactor in serum bottles closed with thick butyl rubber stoppers (#CLS-4209-14, Chemglass Life Sciences) to prevent gas diffusion (only for strains LL345 and LL1715). Gas was flowing through the bottles continuously from the start of the cultivations. These bottles were connected to the top of the condensers *via* septa, sterile 0.6 mm needles, 0.2 µm syringe filters (PES) and Norprene tubing (3 mm ID). The bottles were detached immediately prior to liquid sampling and stored at room temperature until analysis on the same or next day.

The mass spectrometer consisted of an Extrel RGA mounted on an ultrahigh vacuum (<10^-9^ mbar) chamber. Sample gases were injected through a blue butyl rubber stopper (#CLS-4209-14, Chemglass Life Sciences) into a small high-pressure region (pumped between samples) from where the gas was let into the UHV region with a precision leak valve. The pressure in the UHV chamber was monitored and controlled during measurements, typically at 4.5 ± 0.8 · 10^-7^ mbar. Twenty mass spectra were averaged to reduce the effects of any possible changes in pressure during each measurement. Mass spectra for background subtraction (ten) were measured prior to each sample.

Ions were detected using a Faraday cup. The different ionization efficiencies, fragmentation of the different species, and the m/z-dependent transmission efficiency of the quadrupole was accounted for by calibration with gas mixtures of known composition (0.2%, 0.5% and 2% H_2_ in 20% O_2_ and balance N_2_; 1% H_2_ in 99% N_2_ [Skandinaviska Gasprodukter AB, Södertälje, Sweden]; 5% H_2_ in 10% CO_2_ and 85% N_2_ [Strandmöllen, Ljungby, Sweden]) by determining calibration values for each of the parent ions that yielded the correct, known composition. This simple method was only valid for the species in the calibration mixtures but sufficed, as other species (e.g. ethanol) were not observed in the mass spectra of the samples.

### Extracellular metabolite analysis

A culture sample of approximately 100 mL was quickly withdrawn from the reactor and stored on ice. Samples from the feed vessels were collected at the beginning and end of the cultivation. Aliquots of 1 mL were centrifuged in a table-top centrifuge (Eppendorf 5424, Thermo Fisher Scientific, Stockholm, Sweden) at 20,238 *x g* for 2 min. The supernatant was transferred to a nylon, low protein-binding, 0.22-µm Corning Costar Spin-X centrifuge tube filter (Sigma-Aldrich) and centrifuged at 20,238 *x g* for 2 min. Part of the filtrate was stored at 4°C until analysis of sugars and canonical fermentation products on the HPLC, whereas the other part was stored at -20°C until analysis of ammonium, proteins and amino acids (described below).

Supernatant samples stored at 4°C were analyzed within one week for cellobiose, glucose, ethanol, acetate, formate, lactate, pyruvate and malate on a Waters Alliance 2695 HPLC (Waters Sverige AB, Solna, Sweden) equipped with a Bio-Rad Aminex HPX-87H column (Bio-Rad, Solna, Sweden) operated at 60°C and with a 0.6 mL min^-1^ flow rate. Formate, pyruvate, and malate were separated using a 75 mM H_2_SO_4_ mobile phase, whereas remaining components were separated using a 5 mM H_2_SO_4_ mobile phase. Cellobiose, glucose and ethanol were detected by a Waters 2414 refractive-index detector and the acids were detected by a Waters 2996 photodiode-array detector at 210 nm.

Amino acids were analyzed by derivatization using a Waters AccQ-Tag Ultra commercial kit (cat. no. WAT052880) (Waters Sverige AB, Solna, Sweden). First, 2-aminobutyric acid (AABA) (Sigma-Aldrich) was added as an internal standard. Then, supernatant protein was removed by acidification with 5 g L^-1^ trichloroacetic acid (Supelco from Sigma-Aldrich), incubation for 20 min in room temperature and centrifugation at 20,238 *x g* for 10 min in a table-top centrifuge (Eppendorf 5424, Thermo Fisher Scientific, Stockholm, Sweden). The supernatant was derivatized according to the AccQ-Tag commercial kit protocol (Waters “UPLC Amino Acid Analysis Solution System Guide”, 71500129702, revision B). Separation was performed in Waters Acquity Ultra-Performance Liquid Chromatography (UPLC) with a reverse phase-UPLC AccQ-Tag Ultra column (cat.no. WAT052885) at 55°C, with a flow rate of 0.70 mL min^-1^ and a run time of 10 min. The mobile phase consisted of eluent A, prepared by diluting Waters AccQ-Tag Ultra Eluent A concentrate (cat. no. 186003838) twenty times in ultrapure water (Purelab Chorus from AB Ninolab [Stockholm, Sweden]), and eluent B as provided by Waters (cat. no. 186003839). The following elution profile was used: initial 99.9% A, 0.1% B; 0.54 min 99.9% A, 0.1% B, curve 6; 3.80 min, 96.0% A, 4.0% B, curve 6; 4.20 min, 96.0% A, 4.0% D, curve 6; 5.74 min, 90.9% A, 9.1% B, curve 7; 7.74 min, 78.8% A, 21.2% B, curve 6; 8.04 min 40.4% A, 59.6% B, curve 6; 8.05 min, 10.0% A, 90.0 % B, curve 6; 8.64 min, 10.0% A, 90.0% B, curve 6; 8.73 min, 99.9% A, 0.1 % B, curve 6; 9.5 min, 99.9% A, 0.1% B, curve 6. Tryptophan, cysteine, and tyrosine were quantified using an Acquity TUV detector at 250 nm. Alanine, arginine, asparagine, aspartate, glutamine, glycine, glutamate, histidine, isoleucine, leucine, methionine, phenylalanine, serine, threonine, tryptophan, tyrosine, and valine were detected using a Acquity FLR detector with excitation at 266 nm and emission at 473 nm. The UPLC was set-up according to the AccQ-Tag commercial kit protocol. As standards, Waters amino acid standard solution with 17 amino acids (cat. no. WAT088122) was mixed with glutamine (Irvine Scientific, Tilburg, The Netherlands), asparagine, and tryptophan (Sigma-Aldrich). Co-elution of arginine and glutamine was observed.

Extracellular ammonium in the culture supernatants and medium samples was measured spectrophotometrically using a commercial MegaZyme Ammonium Assay Kit (cat. no. KURAMR, Megazyme, Bray, Ireland) and a standard curve with ammonium chloride (Sigma-Aldrich).

### Extracellular protein analysis

Extracellular supernatant protein was quantified using a Bradford assay (56) modified for detection of lower protein concentrations (1-10 µg mL^-1^). The protocol described in Sigma-Aldrich technical bulletin (cat. no. B6916) was used with a 96-well microplate (flat-bottom, clear, PS) (Sigma-Aldrich) and bovine serum albumin (Sigma-Aldrich) as standard. A conversion to molar carbon, nitrogen and degree-of-reduction was done using an estimated general amino acid composition for *C. thermocellum*, which was derived by counting the occurrences of each amino acid in all open-reading frames of the genome and correcting for polymerization (loss of H_2_O) (see Supplemental Table S1). These elemental concentrations were used to calculate respective recovery on substrate (described below) (Supplemental Table S3-S5).

### CDW and OD measurements

Cell dry weight (CDW) was quantified in technical triplicates by first transferring 10 mL of culture into dried, pre-weighed, conical glass centrifuge tubes and centrifuging at 2,250 *x g* in a table-top centrifuge (Z206 A, Hermle Labortechnik GmbH, Wehingen, Germany) for 20 min. After washing the pellet in an equal volume of ultrapure water (Purelab Chorus), another round of centrifugation was performed. The washed pellet was dried overnight in a VENTI-Line forced-convention oven (VWR) set at 105°C. Upon cooling for one hour in a desiccator, the dried glass tube was weighed. The CDW was calculated by dividing the difference in tube weights with the sample volume. Optical density (OD) was measured in technical triplicates in 1-mL polystyrene cuvettes in a V-1200 spectrophotometer (VWR) at 600 nm. Deionized water was used as a blank and for dilutions.

### Glycogen determination

The sampling and assay of glycogen content was performed as previously described (25). Briefly, 1 mL of sample was mixed with 5 mL of ice-cold methanol (−80°C) and centrifuged (10,000 *x g*, 10 min, -10°C). The pellet was washed with 5 mL of methanol (−80°C) and centrifuged again. After removing the supernatant, the pellet was stored at −80°C until analysis. Glycogen was hydrolyzed to glucose and analyzed on HPLC (as described above).

### Calculation of yields and rates

Quantification of extracellular metabolites and biomass concentration in chemostats allowed calculation of yields on substrate (Y_i/S_) and biomass-specific rates (q) at steady state. A general mass balance for a metabolite *i* in the liquid phase is described by equation (1).

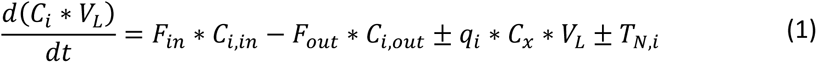

Where *C* denotes concentrations (mmol L^-1^) for metabolite *i* or biomass *x*, *V*_L_is the liquid volume (L), *F* is the flow rate (L h^-1^), *q_i_* is the biomass-specific conversion rate (mmol g_x_^-1^ h^-1^) and *T*_N,i_ is the transfer rate between the gas and liquid phase for volatile compounds (mmol g_x_^-1^ h^-1^). Both *q*_i_ and *T*_N,i_ are defined as positive numbers and the sign in front of each term depends on production/transfer into liquid (+) or consumption/transfer out of liquid (-). Assuming steady state 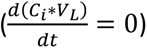, ideal mixing (*C*_i,out_ = *C*_i_), a constant volume, *F* = *F_in_* = *F_out_*, and defining the dilution rate as 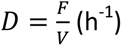, an expression for *q_i_* can be derived.

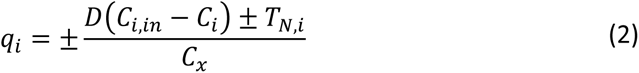

With a condenser to minimize evaporation, the transfer rate, *T_N,i_*, was assumed to be zero for all aqueous metabolites except ethanol. For ethanol, a first-order transfer rate coefficient was determined to *k_ethanol_*= 0.018 ± 0.01 h^-1^ (*n* = 3) in the specified bioreactor system at 55°C, 400 rpm, and 0.2 L min^-1^ overhead purging with N_2_ gas, by measuring dissipating ethanol concentrations in the liquid phase over time (data not shown). Therefore, the transfer term in (2) was −*k_ethanol_* ∗ *C_ethanol_* ∗ *V_L_*, giving following expression for the ethanol production rate (where *C_ethanol,in_* = 0).

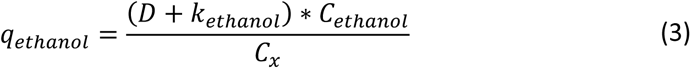

Based on the biomass-specific conversion rates, the yield of product, *i*, on the consumed substrate cellobiose, *S*, was calculated as following.

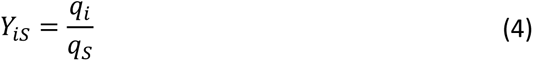

In order to calculate the molar yield of biomass on cellobiose, a molecular weight of 24.66 g Cmol^-1^ and a degree-of-reduction of 4.30 was used based on the biomass composition of CH_1.71_O_0.43_N_0.20_S_0.01_ with 4.32% ash content from Hogsett (57) and following elemental degree of reductions: C = 4, H = 1, O = -2, N = -3, S = 6.

The estimated rate of CO_2_ production was calculated based on current biochemical knowledge on the *C. thermocellum* metabolism and the genome-scale metabolic model iCBI655 (11). In addition, in the conversion of cellobiose into biomass, CO_2_ is produced as a byproduct to balance the carbon since biomass is slightly more reduced per carbon than cellobiose. Based on carbon and degree-of-reduction balancing of that biosynthesis reaction, it was estimated that 3.04 mmol CO_2_ per g biomass is formed. With this, the total CO_2_ production rate was calculated based on measured conversion rates as shown in equation (5) below.

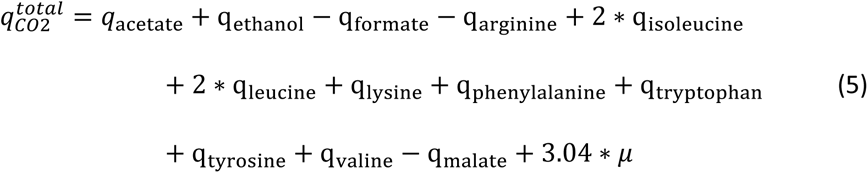

The H_2_ production rate was derived by combining the mass balances for the gas and liquid phase. First, the general mass balance in the gas phase, as shown in equation (6), can be simplified for H_2_ since the inlet gas only consists of N_2_. In steady state, equation (7) is valid.

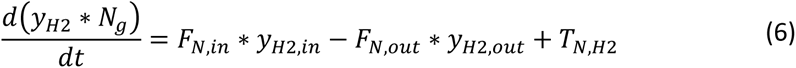

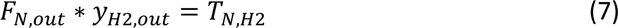

Where *y*_i_ denotes the mole fraction of the gas, *N_g_* is the total gas amount (mol) and *F_N_* is the gas flow in or out of the bioreactor (mol h^-1^). By combining equation (1) for H_2_ (simplified by neglecting in- and outflow of H_2_ in the liquid due to low solubility) with equation (7), the H_2_ production rate was derived.

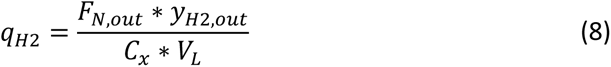

Where *F_N,out_*(mol h^-1^) could be estimated from a nitrogen balance on the inlet (pure N_2_) and outlet (N_2_, CO_2_ and H_2_) gas streams (N_2_ is inert) and using the ideal gas law.

### Calculation of carbon-, nitrogen- and degree-of-reduction recoveries on consumed substrate

The recovery on substrate was defined as the ratio between products and substrates in regard to their carbon, nitrogen or degree-of-reduction content. The substrate for carbon and degree-of-reduction calculations was cellobiose and for nitrogen calculations was ammonium. In the degree-of-reduction balancing, following definitions were made for the elements: C = 4, H = 1, O = -2, N = -3, S = 6. Molar concentrations of biomass and protein were estimated based on molecular weights specified above.

### Enzyme activity assays

Cell-free extracts were prepared as described previously (53). In the case of steady-state cultures, 50 mL sample was harvested. Enzyme activities were assayed aerobically at 55°C in a Cary 50 UV-visible spectrophotometer with a single-cell Peltier element from Varian AB (Solna, Sweden). The conversion of NAD(P)^+^ to NAD(P)H was followed at 340 nm in quartz cuvettes (VWR) with a path length of 1 cm and reaction volume of 1 mL. An extinction coefficient of 6.22 AU mmol^-1^ L cm^-1^ for NAD(P)H was used. The proportionality between the activity and the amount of cell-free extract was confirmed by assaying two different concentrations of cell-free extract in technical duplicates. Each activity is reported for biological duplicates. An assay was carried out by first incubating all reaction mixture components except coupling enzymes and cell-free extract at 55°C for 5 min. Then, coupling enzymes (if used) and cell-free extract was added and equilibrated for an additional 4 minutes, with the last minute serving as background slope. The reaction was initiated by adding indicated metabolites. The slope for the first 30 seconds was used to calculate the activity. Glutamate dehydrogenase activity was measured based on the oxidation rate of NAD(P)H as 2-ketoglutarate is converted to glutamate (10, 58). The assay mixture contained 50 mM Tris-HCl (pH 8.0 at 55°C), 5 mM DTT, 50 mM NH_4_Cl, 5 mM 2-ketoglutarate (pH 8.0 at 25°C), and 0.3 mM NADPH or NADH. The reaction was started by addition of 2-ketoglutarate.

Malic enzyme activity was measured by following the reduction of NADP^+^ as malate is converted to pyruvate (26). The assay mixture contained 50 mM Tris-HCl (pH 7.5 at 55°C), 5 mM DTT, 2 mM NADP^+^, 20 mM NH_4_Cl, 2.5 mM malate, and 50 or 100 µL cell-free extract. The reaction was started by addition of malate.

Pyruvate phosphate dikinase activity was assayed by coupling the formation of pyruvate from PEP to the formation of lactate from pyruvate via added lactate dehydrogenase, resulting in the oxidation of NADH (26). The assay mixture contained 50 mM Tris-HCl (pH 7.5 at 55°C), 5 mM DTT, 0.3 mM NADH, 5 mM MgCl_2_·6H_2_O, 20 mM NH_4_Cl, 2 mM PEP, 2 mM AMP, 13 U mL^-1^ L-lactate dehydrogenase (from bovine heart, Sigma L2625), 1 mM K_4_PP_i_, and 50 or 100 µL cell-free extract. The reaction was started by addition of PP_i_.

Pyruvate kinase activity was measured by coupling the formation of pyruvate from PEP to the formation of lactate from pyruvate via added lactate dehydrogenase, resulting in oxidation of NADH (26). The assay mixture contained 50 mM Tris-HCl (pH 7.5 at 55°C), 5 mM DTT, 0.3 mM NADH, 12 mM MgCl_2_·6H_2_O, 10 mM KCl, 10 mM ADP, 0.1 mM 3-phosphoglyceric acid, 5 mM PEP, 13 U mL^-1^ L-lactate dehydrogenase (from bovine heart, Sigma L2625), and 50 or 100 µL cell-free extract. The reaction was initiated by adding PEP.

Lactate dehydrogenase (LDH) activity was assayed by measuring oxidation of NADH as pyruvate is converted to lactate (59). The mixture contained 10 mM pyruvate, 1 mM fructose 1,6-bisphosphate, 0.22 mM NADH, 200 mM Tris-HCl (pH 7.3 at 55°C), and 50 or 100 ml of cell-free extract. The reaction was initiated by adding pyruvate. LDH activity was used routinely as quality control on all cell-free extracts (see Supplemental Table S8).

### Data analysis

Student’s *t* test was used for unpaired comparison between values in this study. **Data availability.** Data on measured concentrations from the chemostats to calculate yields and rates are presented in Supplemental Table S7. Genbank accession numbers for plasmids generated in this study are in Table 6.

## ACKNOWLEDGEMENTS

Funding for J.Y., T.K., A.K., and A.J.A.M was provided by Formas grant 2017-00973 and the Novo Nordisk Foundation grant NNF20OC0064164. D.J.H. was funded by the Swedish Foundation for Strategic Research (SSF) (ITM17-0236). T.R. and A.M.G. were supported by the Center for Bioenergy Innovation, U.S. DOE Bioenergy Research Center, supported by the Office of Biological and Environmental Research in the DOE Office of Science. Oak Ridge National Laboratory is managed by UT-Battelle, LLC, for the U.S. DOE under contract DE-AC05-00OR22725.

We thank Gustav Sjöberg, Jeroen G. Koendjbiharie and Chonticha Phongsawat for insightful comments on the data analysis as well as for help in setting up and sampling the chemostats. We are also thankful to Amparo Jiménez Quero for experimental help with metabolite analysis.

## SUPPLEMENTAL TABLES

**TABLE S1.**
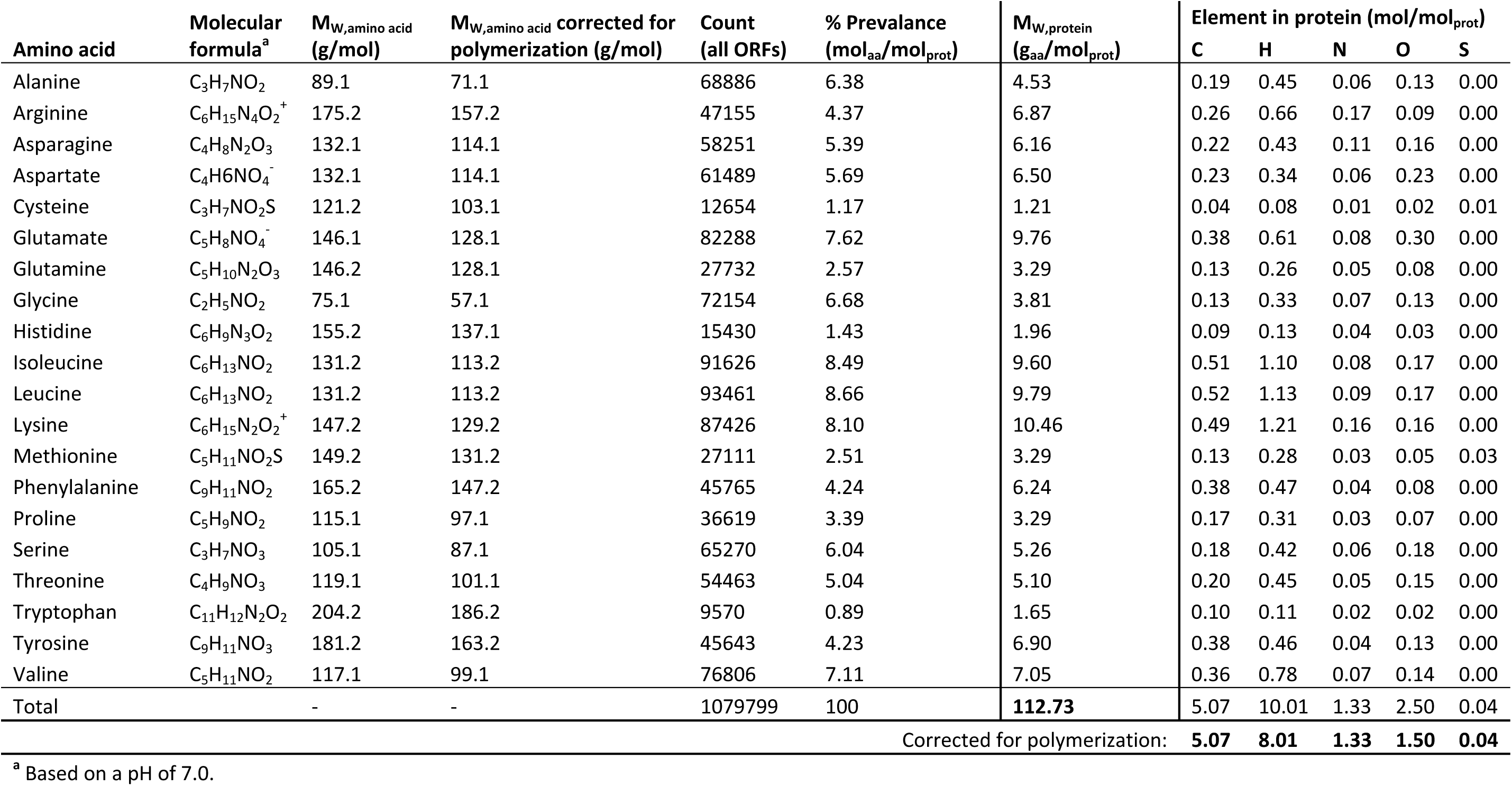
Molecular weight and elemental composition of a general protein in the C. thermocellum ATCC 27405 genome by counting the urrences of each amino acid in all open-reading frames and correcting for polymerization (loss of H_2_O).

**TABLE S2.** Amino acid yields (mmol mol_cellobiose_^-1^) in chemostats with either cellobiose or ammonium as the sole limiting nutrient at a dilution e of 0.10 h^-1^.^a^

|  | Cellobiose-limitation |  |  |  |  | Ammonium-limitation |  |  |  |  |
| --- | --- | --- | --- | --- | --- | --- | --- | --- | --- | --- |
|  | Reference strain (LL345) | AVM064 (*Δppdk ΔmalE::P<sub>eno</sub>-pyk*) | AVM003 (*Δppdk*) | LL1084 (*ΔnfnAB*) | AG1715 (*Δgogat*) | Reference strain (LL345) | AVM064 (*Δppdk ΔmalE::P<sub>eno</sub>-pyk*) | AVM003 (*Δppdk*) | LL1084 (*ΔnfnAB*) | AG1715 (*Δgogat*) |
| Alanine | 6.5 ± 0.2 | 6.7 ± 0.4 | 6.7 ± 0.5 | 6.8 ± 0.2 | 5.6 ± 0.1 | 9.5 ± 0.1 | 6.1 ± 0.4 | 9.0 ± 0.4 | 8.0 ± 0.4 | 5.3 ± 0.4 |
| Asparagine | <0.5 | <0.5 | <0.5 | <0.5 | <0.5 | <0.5 | <0.5 | <0.5 | <0.5 | <0.5 |
| Aspartate | <0.5 | 2.2 ± 0.3 | 1.0 ± 0.1 | 0.9 ± 0.5 | 1.1 ± 0.7 | <0.5 | 0.8 ± 0.2 | <0.5 | 0.7 ± 0.1 | <0.5 |
| Glycine | 2.0 ± 0.2 | 2.4 ± 0.2 | 2.2 ± 0.2 | 1.9 ± 0.0 | 1.5 ± 0.2 | 2.2 ± 0.1 | 1.6 ± 0.2 | 1.9 ± 0.2 | 2.0 ± 0.2 | <0.5 |
| Glutamate | 2.8 ± 0.1 | 2.7 ± 0.4 | 2.2 ± 0.1 | 3.1 ± 0.2 | 2.8 ± 0.2 | 5.8 ± 0.1 | 4.2 ± 0.2 | 4.9 ± 0.2 | 8.3 ± 0.4 | 5.5 ± 0.5 |
| Glutamine<sup>b</sup> | <0.7 ± 0.0 | <2.0 ± 0.4 | <2.1 ± 0.7 | <1.7 ± 0.2 | <0.6 ± 0.1 | <0.5 | <2.1 ± 0.2 | <2.8 ± 0.3 | <1.0 ± 0.1 | <0.5 |
| Histidine | <0.5 | <0.5 | <0.5 | <0.5 | <0.5 | <0.5 | <0.5 | <0.5 | <0.5 | <0.5 |
| Isoleucine | 1.3 ± 0.0 | 0.7 ± 0.0 | 1.5 ± 0.1 | 1.6 ± 0.2 | 1.1 ± 0.0 | 10.0 ± 0.2 | 3.7 ± 0.4 | 11.2 ± 2.4 | 9.1 ± 0.8 | 2.5 ± 0.8 |
| Leucine | 0.8 ± 0.0 | 1.4 ± 0.1 | <0.5 | 0.7 ± 0.6 | 0.8 ± 0.0 | 2.5 ± 0.1 | 2.3 ± 0.2 | 2.9 ± 0.1 | 2.8 ± 0.1 | 1.1 ± 0.2 |
| Methionine | <0.5 | <0.5 | <0.5 | <0.5 | <0.5 | 6.3 ± 0.5 | <0.5 | <0.5 | 0.9 ± 0.3 | <0.5 |
| Phenylalanine | 1.1 ± 0.1 | 4.2 ± 0.6 | 1.2 ± 0.3 | 0.9 ± 0.2 | 0.9 ± 0.1 | 4.2 ± 0.1 | 4.4 ± 1.3 | 4.2 ± 0.5 | 5.8 ± 0.2 | 4.7 ± 1.0 |
| Serine | 2.6 ± 0.2 | 2.1 ± 0.1 | 2.3 ± 0.0 | 2.5 ± 0.1 | 2.0 ± 0.1 | 1.9 ± 0.2 | 1.4 ± 0.1 | 2.0 ± 0.1 | 2.0 ± 0.2 | 1.2 ± 0.5 |
| Threonine | 3.4 ± 0.3 | 3.8 ± 0.4 | 3.5 ± 0.4 | 3.6 ± 0.2 | 2.9 ± 0.3 | 3.6 ± 0.1 | 3.6 ± 0.4 | 3.0 ± 0.4 | 5.2 ± 0.5 | <0.5 |
| Tryptophan | <0.5 | <0.5 | <0.5 | <0.5 | <0.5 | <0.5 | <0.5 | <0.5 | <0.5 | <0.5 |
| Tyrosine | 0.9 ± 0.0 | 0.8 ± 0.1 | <0.5 | <0.5 | <0.5 | 1.9 ± 0.1 | 1.1 ± 0.1 | 1.4 ± 0.1 | 4.0 ± 1.0 | <0.5 |
| Valine | 3.3 ± 0.1 | 1.3 ± 0.1 | 5.1 ± 0.3 | 5.8 ± 0.3 | 2.3 ± 0.1 | 67.3 ± 1.7 | 12.6 ± 2.8 | 62.3 ± 23.1 | 65.3 ± 8.3 | 16.0 ± 9.4 |

<sup>a</sup> Data are shown as average ± standard deviation for three biological replicates
<sup>b</sup> Estimated upper limit (see Materials and Methods)

**Table S3.** Carbon recovery on consumed cellobiose in chemostats limited on either cellobiose or ammonium at a dilution rate of 0.10 h^-1^.^a^

| % (Cmol/Cmol) | Cellobiose-limitation |  |  |  |  | Ammonium-limitation |  |  |  |  |
| --- | --- | --- | --- | --- | --- | --- | --- | --- | --- | --- |
|  | Reference strain (LL345) | AVM064 (*Δppdk ΔmalE::P<sub>eno</sub>-pyk*) | AVM003 (*Δppdk*) | LL1084 (*ΔnfnAB*) | AG1715 (*Δgogat*) | Reference strain (LL345) | AVM064 (*Δppdk ΔmalE::P<sub>eno</sub>-pyk*) | AVM003 (*Δppdk*) | LL1084 (*ΔnfnAB*) | AG1715 (*Δgogat*) |
| Biomass<sup>b</sup> | 15.49 ± 0.27 | 14.34 ± 0.61 | 16.79 ± 0.76 | 15.07 ± 0.04 | 15.79 ± 0.20 | 13.48 ± 0.12 | 13.21 ± 0.51 | 14.60 ± 0.64 | 13.49 ± 0.62 | 16.03 ± 0.57 |
| Glucose | 0.16 ± 0.02 | 0.06 ± 0.02 | 0.07 ± 0.03 | 0.03 ± 0.00 | 0.08 ± 0.03 | 0.59 ± 0.07 | 1.02 ± 0.18 | 1.56 ± 0.20 | 1.39 ± 0.12 | 1.55 ± 0.38 |
| Ethanol | 16.16 ± 0.97 | 25.78 ± 0.17 | 16.00 ± 0.55 | 16.04 ± 0.27 | 13.99 ± 1.10 | 17.76 ± 0.46 | 25.86 ± 0.89 | 14.06 ± 0.82 | 16.73 ± 0.42 | 12.91 ± 0.56 |
| Acetate | 23.04 ± 0.22 | 10.81 ± 0.30 | 24.29 ± 0.72 | 23.01 ± 0.31 | 23.00 ± 0.25 | 15.60 ± 0.10 | 10.09 ± 0.17 | 19.57 ± 2.00 | 16.85 ± 0.22 | 14.00 ± 1.13 |
| Formate | 0.96 ± 0.31 | 1.87 ± 0.16 | 0.81 ± 0.10 | 2.56 ± 0.06 | 0.62 ± 0.06 | 1.23 ± 0.09 | 2.64 ± 0.27 | 1.52 ± 0.43 | 1.04 ± 0.23 | 2.48 ± 1.02 |
| Lactate | 0.35 ± 0.03 | 0.51 ± 0.07 | 0.43 ± 0.05 | 0.30 ± 0.05 | 0.40 ± 0.05 | 1.68 ± 0.14 | 1.02 ± 0.41 | 1.53 ± 0.51 | 1.41 ± 0.10 | 0.92 ± 0.45 |
| Pyruvate | 0.09 ± 0.01 | 0.06 ± 0.02 | 0.33 ± 0.21 | 0.21 ± 0.04 | 0.27 ± 0.02 | 2.25 ± 0.12 | 0.28 ± 0.05 | 0.57 ± 0.11 | 0.77 ± 0.10 | 0.58 ± 0.29 |
| Malate | 0.16 ± 0.16 | 3.64 ± 0.62 | <0.01 | <0.01 | <0.01 | <0.01 | 1.13 ± 0.11 | <0.01 | 0.07 ± 0.04 | <0.01 |
| CO<sub>2</sub><sup>c</sup> | 19.83 ± 0.16 | 16.68 ± 0.36 | 20.66 ± 0.48 | 18.19 ± 0.26 | 19.11 ± 0.42 | 17.26 ± 0.23 | 16.29 ± 0.07 | 17.18 ± 0.69 | 17.55 ± 0.18 | 12.36 ± 0.97 |
| H<sub>2</sub> | 0 | 0 | 0 | 0 | 0 | 0 | 0 | 0 | 0 | 0 |
| Ex. protein<sup>d</sup> | 3.94 ± 0.11 | 2.36 ± 0.18 | 3.19 ± 0.18 | 4.08 ± 0.14 | 3.40 ± 0.08 | 2.00 ± 0.08 | 1.50 ± 0.16 | 2.29 ± 0.14 | 1.87 ± 0.06 | 1.86 ± 0.25 |
| Alanine | 0.16 ± 0.01 | 0.17 ± 0.01 | 0.17 ± 0.01 | 0.17 ± 0.01 | 0.14 ± 0.00 | 0.24 ± 0.00 | 0.15 ± 0.01 | 0.22 ± 0.01 | 0.20 ± 0.01 | 0.13 ± 0.01 |
| Asparagine | <0.01 | <0.01 | <0.01 | <0.01 | <0.01 | <0.01 | <0.01 | <0.01 | <0.01 | <0.01 |
| Aspartate | 0.01 ± 0.02 | 0.07 ± 0.01 | 0.03 ± 0.00 | 0.03 ± 0.02 | 0.04 ± 0.02 | 0.01 ± 0.00 | 0.03 ± 0.01 | <0.01 | 0.02 ± 0.00 | <0.01 |
| Glycine | 0.04 ± 0.01 | 0.04 ± 0.00 | 0.04 ± 0.00 | 0.03 ± 0.00 | 0.03 ± 0.00 | 0.04 ± 0.00 | 0.03 ± 0.00 | 0.03 ± 0.00 | 0.03 ± 0.00 | <0.01 |
| Glutamate | 0.12 ± 0.01 | 0.11 ± 0.02 | 0.09 ± 0.01 | 0.13 ± 0.01 | 0.12 ± 0.01 | 0.24 ± 0.01 | 0.17 ± 0.01 | 0.20 ± 0.01 | 0.35 ± 0.01 | 0.23 ± 0.02 |
| Glutamine/arginine<sup>e</sup> | 0.03 ± 0.00 | 0.08 ± 0.02 | 0.09 ± 0.03 | 0.07 ± 0.01 | 0.02 ± 0.00 | 0.02 ± 0.01 | 0.09 ± 0.01 | 0.12 ± 0.01 | 0.04 ± 0.00 | 0.02 ± 0.00 |
| Histidine | <0.01 | <0.01 | <0.01 | <0.01 | <0.01 | <0.01 | <0.01 | 0.02 ± 0.00 | <0.01 | <0.01 |
| Isoleucine | 0.06 ± 0.00 | 0.03 ± 0.00 | 0.07 ± 0.00 | 0.08 ± 0.01 | 0.05 ± 0.00 | 0.50 ± 0.01 | 0.18 ± 0.02 | 0.56 ± 0.12 | 0.45 ± 0.04 | 0.12 ± 0.04 |
| Leucine | 0.04 ± 0.00 | 0.07 ± 0.01 | <0.01 | 0.03 ± 0.03 | 0.04 ± 0.00 | 0.13 ± 0.01 | 0.11 ± 0.01 | 0.15 ± 0.00 | 0.14 ± 0.01 | 0.06 ± 0.01 |
| Methionine | <0.01 | <0.01 | <0.01 | <0.01 | <0.01 | 0.26 ± 0.02 | <0.01 | 0.02 ± 0.02 | 0.04 ± 0.01 | <0.01 |
| Phenylalanine | 0.08 ± 0.01 | 0.31 ± 0.04 | 0.09 ± 0.02 | 0.07 ± 0.01 | 0.07 ± 0.00 | 0.31 ± 0.01 | 0.33 ± 0.10 | 0.31 ± 0.04 | 0.43 ± 0.02 | 0.35 ± 0.08 |
| Serine | 0.06 ± 0.01 | 0.05 ± 0.00 | 0.06 ± 0.00 | 0.06 ± 0.00 | 0.05 ± 0.00 | 0.05 ± 0.01 | 0.03 ± 0.00 | 0.05 ± 0.00 | 0.05 ± 0.00 | 0.03 ± 0.01 |
| Threonine | 0.11 ± 0.01 | 0.13 ± 0.01 | 0.12 ± 0.01 | 0.12 ± 0.01 | 0.10 ± 0.01 | 0.12 ± 0.00 | 0.12 ± 0.01 | 0.10 ± 0.01 | 0.17 ± 0.02 | 0.02 ± 0.01 |
| Tryptophan | <0.01 | 0.03 ± 0.00 | <0.01 | <0.01 | <0.01 | <0.01 | <0.01 | <0.01 | <0.01 | <0.01 |
| Tyrosine | 0.07 ± 0.00 | 0.06 ± 0.01 | 0.04 ± 0.00 | <0.01 | <0.01 | 0.14 ± 0.01 | 0.08 ± 0.01 | 0.10 ± 0.01 | 0.30 ± 0.08 | <0.01 |
| Valine | 0.14 ± 0.01 | 0.05 ± 0.00 | 0.21 ± 0.01 | 0.24 ± 0.01 | 0.09 ± 0.00 | 2.80 ± 0.07 | 0.53 ± 0.12 | 2.59 ± 0.96 | 2.72 ± 0.35 | 0.67 ± 0.39 |
| Sum of all products | 81 ± 1 | 77 ± 0 | 84 ± 2 | 81 ± 1 | 77 ± 1 | 77 ± 1 | 75 ± 1 | 77 ± 3 | 76 ± 1 | 64 ± 1 |
| Sum of amino acids | 0.93 ± 0.05 | 1.22 ± 0.05 | 1.01 ± 0.07 | 1.04 ± 0.06 | 0.75 ± 0.04 | 4.86 ± 0.07 | 1.88 ± 0.29 | 4.49 ± 1.11 | 4.96 ± 0.34 | 1.64 ± 0.42 |

<sup>a</sup> Data are shown as average ± standard deviation for three biological replicates. <sup>b</sup> Based on a molecular weight of 24.66 g mol<sup>-1</sup> for the biomass composition CH<sub>1.71</sub>O<sub>0.43</sub>N<sub>0.20</sub>S<sub>0.01</sub> with 4.32 % ash content (1). <sup>c</sup> Calculated from fermentation product yields and biomass yield (see Materials and Methods). <sup>d</sup> Based on a molecular weight of a general protein (see Supplemental Table S2 and Materials and Methods). <sup>e</sup> Estimated lower value based on 4 carbons per mole.

**Table S4.**
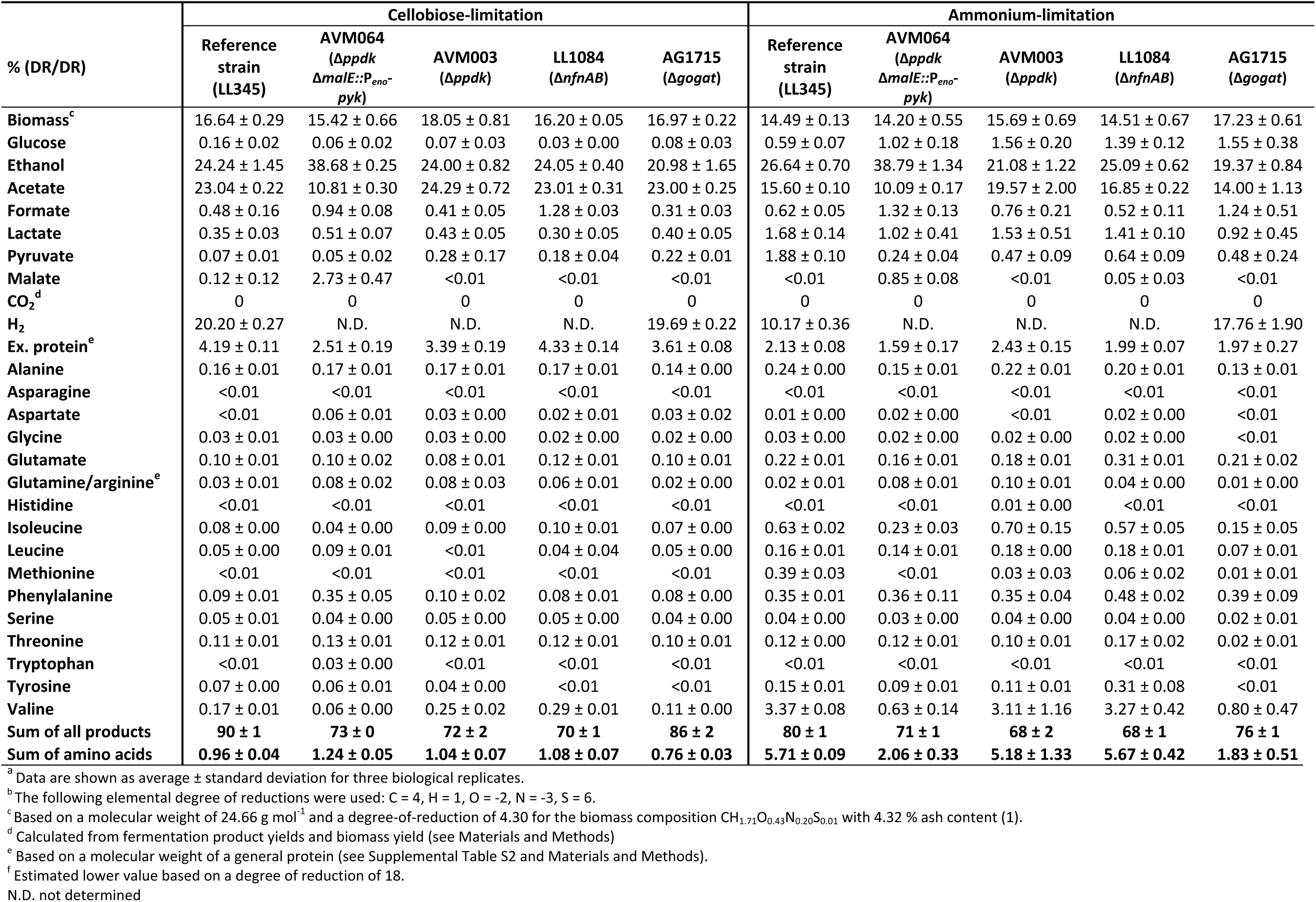
Degree-of-reduction recovery on consumed cellobiose in chemostats limited on either cellobiose or ammonium at 0.10 h^-1^.^a,b^

**Table S5.** Nitrogen recovery on consumed ammonium in chemostats limited on either cellobiose or ammonium at a dilution rate of 0.10 h^-1^.^a^

| % (Nmol/Nmol) | Cellobiose-limitation |  |  |  |  | Ammonium-limitation |  |  |  |  |
| --- | --- | --- | --- | --- | --- | --- | --- | --- | --- | --- |
|  | Reference strain (LL345) | AVM064 (*Δppdk* *ΔmalE::P<sub>eno</sub>-pyk*) | AVM003 (*Δppdk*) | LL1084 (*ΔnfnAB*) | AG1715 (*Δgogat*) | Reference strain (LL345) | AVM064 (*Δppdk* *ΔmalE::P<sub>eno</sub>-pyk*) | AVM003 (*Δppdk*) | LL1084 (*ΔnfnAB*) | AG1715 (*Δgogat*) |
| Biomass<sup>b</sup> | 50.45 ± 1.13 | 49.96 ± 1.52 | 53.18 ± 1.83 | 37.10 ± 2.43 | 39.18 ± 1.10 | 56.01 ± 1.61 | 72.80 ± 3.19 | 62.93 ± 4.45 | 62.60 ± 1.72 | 57.36 ± 4.53 |
| Glucose | 0 | 0 | 0 | 0 | 0 | 0 | 0 | 0 | 0 | 0 |
| Ethanol | 0 | 0 | 0 | 0 | 0 | 0 | 0 | 0 | 0 | 0 |
| Acetate | 0 | 0 | 0 | 0 | 0 | 0 | 0 | 0 | 0 | 0 |
| Formate | 0 | 0 | 0 | 0 | 0 | 0 | 0 | 0 | 0 | 0 |
| Lactate | 0 | 0 | 0 | 0 | 0 | 0 | 0 | 0 | 0 | 0 |
| Pyruvate | 0 | 0 | 0 | 0 | 0 | 0 | 0 | 0 | 0 | 0 |
| Malate | 0 | 0 | 0 | 0 | 0 | 0 | 0 | 0 | 0 | 0 |
| CO<sub>2</sub><sup>c</sup> | 0 | 0 | 0 | 0 | 0 | 0 | 0 | 0 | 0 | 0 |
| H<sub>2</sub> | 0 | 0 | 0 | 0 | 0 | 0 | 0 | 0 | 0 | 0 |
| Ex. protein<sup>d</sup> | 16.95 ± 1.09 | 10.85 ± 0.48 | 13.03 ± 0.43 | 13.21 ± 0.59 | 11.14 ± 0.56 | 10.97 ± 0.44 | 10.89 ± 1.08 | 13.01 ± 1.02 | 11.47 ± 0.50 | 8.71 ± 0.56 |
| Alanine | 0.89 ± 0.05 | 0.98 ± 0.12 | 0.93 ± 0.03 | 0.70 ± 0.06 | 0.58 ± 0.03 | 1.66 ± 0.06 | 1.41 ± 0.10 | 1.62 ± 0.07 | 1.56 ± 0.04 | 0.80 ± 0.13 |
| Asparagine | 0.04 ± 0.00 | 0.07 ± 0.01 | 0.05 ± 0.00 | 0.01 ± 0.00 | 0.01 ± 0.00 | 0.02 ± 0.03 | 0.03 ± 0.00 | 0.02 ± 0.00 | 0.03 ± 0.00 | <0.01 |
| Aspartate | 0.04 ± 0.06 | 0.32 ± 0.04 | 0.14 ± 0.01 | 0.09 ± 0.05 | 0.11 ± 0.07 | 0.06 ± 0.01 | 0.18 ± 0.04 | 0.05 ± 0.00 | 0.13 ± 0.02 | 0.02 ± 0.01 |
| Glycine | 0.28 ± 0.02 | 0.35 ± 0.04 | 0.28 ± 0.03 | 0.19 ± 0.02 | 0.16 ± 0.02 | 0.38 ± 0.01 | 0.38 ± 0.05 | 0.35 ± 0.02 | 0.38 ± 0.03 | 0.06 ± 0.02 |
| Glutamate | 0.38 ± 0.02 | 0.39 ± 0.05 | 0.29 ± 0.01 | 0.32 ± 0.04 | 0.29 ± 0.02 | 1.01 ± 0.05 | 0.96 ± 0.05 | 0.88 ± 0.04 | 1.62 ± 0.06 | 0.83 ± 0.12 |
| Glutamine/arginine<sup>e</sup> | 0.19 ± 0.01 | 0.60 ± 0.16 | 0.45 ± 0.03 | 0.35 ± 0.04 | 0.12 ± 0.02 | 0.16 ± 0.07 | 0.98 ± 0.10 | 1.00 ± 0.09 | 0.38 ± 0.04 | 0.11 ± 0.02 |
| Histidine | <0.01 | <0.01 | <0.01 | <0.01 | <0.01 | 0.01 ± 0.00 | 0.12 ± 0.03 | 0.19 ± 0.05 | 0.08 ± 0.01 | 0.05 ± 0.05 |
| Isoleucine | 0.17 ± 0.01 | 0.10 ± 0.01 | 0.20 ± 0.01 | 0.17 ± 0.03 | 0.11 ± 0.00 | 1.74 ± 0.02 | 0.85 ± 0.11 | 2.02 ± 0.39 | 1.76 ± 0.14 | 0.38 ± 0.16 |
| Leucine | 0.11 ± 0.01 | 0.21 ± 0.03 | <0.01 | 0.07 ± 0.06 | 0.08 ± 0.00 | 0.44 ± 0.01 | 0.53 ± 0.04 | 0.52 ± 0.01 | 0.55 ± 0.01 | 0.17 ± 0.04 |
| Methionine | <0.01 | <0.01 | <0.01 | <0.01 | 0.01 ± 0.02 | 1.09 ± 0.10 | 0.03 ± 0.01 | 0.07 ± 0.09 | 0.17 ± 0.05 | 0.04 ± 0.03 |
| Phenylalanine | 0.15 ± 0.01 | 0.61 ± 0.05 | 0.14 ± 0.01 | 0.09 ± 0.02 | 0.09 ± 0.01 | 0.72 ± 0.04 | 1.01 ± 0.33 | 0.75 ± 0.11 | 1.12 ± 0.06 | 0.69 ± 0.14 |
| Serine | 0.35 ± 0.04 | 0.30 ± 0.04 | 0.30 ± 0.00 | 0.26 ± 0.02 | 0.21 ± 0.01 | 0.33 ± 0.03 | 0.32 ± 0.01 | 0.35 ± 0.03 | 0.39 ± 0.04 | 0.17 ± 0.06 |
| Threonine | 0.46 ± 0.03 | 0.56 ± 0.07 | 0.45 ± 0.04 | 0.37 ± 0.04 | 0.30 ± 0.04 | 0.63 ± 0.01 | 0.83 ± 0.10 | 0.54 ± 0.05 | 1.01 ± 0.08 | 0.07 ± 0.06 |
| Tryptophan | 0.01 ± 0.01 | 0.09 ± 0.01 | 0.03 ± 0.00 | <0.01 | <0.01 | <0.01 | 0.02 ± 0.00 | 0.02 ± 0.00 | <0.01 | <0.01 |
| Tyrosine | 0.13 ± 0.01 | 0.12 ± 0.01 | 0.07 ± 0.00 | <0.01 | <0.01 | 0.33 ± 0.02 | 0.26 ± 0.03 | 0.25 ± 0.02 | 0.77 ± 0.21 | <0.01 |
| Valine | 0.45 ± 0.01 | 0.19 ± 0.02 | 0.66 ± 0.03 | 0.59 ± 0.04 | 0.24 ± 0.01 | 11.71 ± 0.11 | 2.92 ± 0.68 | 11.17 ± 3.86 | 12.69 ± 1.43 | 2.48 ± 1.68 |
| Sum of all products | 71 ± 2 | 66 ± 2 | 47 ± 41 | 54 ± 3 | 53 ± 2 | 87 ± 2 | 95 ± 3 | 96 ± 2 | 97 ± 1 | 72 ± 6 |
| Sum of amino acids | 3.66 ± 0.00 | 4.88 ± 0.49 | 2.66 ± 2.31 | 3.23 ± 0.36 | 2.31 ± 0.21 | 20.28 ± 0.36 | 10.83 ± 1.55 | 19.79 ± 4.23 | 22.65 ± 1.40 | 5.87 ± 2.06 |

<sup>a</sup> Data are shown as average ± standard deviation for three biological replicates. <sup>b</sup> Based on a molecular weight of 24.66 g mol<sup>-1</sup> for the biomass composition CH<sub>1.71</sub>O<sub>0.43</sub>N<sub>0.26</sub>S<sub>0.01</sub> with 4.32 % ash content (1). <sup>c</sup> Calculated from fermentation product yields and biomass yield (see Materials and Methods) <sup>d</sup> Based on a molecular weight of a general protein (see Supplemental Table S2 and Materials and Methods). <sup>e</sup> Estimated lower value based on 2 nitrogens per mole.

**TABLE S6.** Biomass-specific conversion rates in chemostats limited on either cellobiose or ammonium at a dilution rate of 0.10 h^-1^.^a,b^

| Unit: mmol g<sub>biomass</sub><sup>-1</sup> h<sup>-1</sup> | Cellobiose-limitation |  |  |  |  | Ammonium-limitation |  |  |  |  |
| --- | --- | --- | --- | --- | --- | --- | --- | --- | --- | --- |
| | Reference strain (LL345) | AVM064 (Δppdk ΔmalE::P<sub>eno</sub><sup>-</sup> pyk) | AVM003 (Δppdk) | LL1084 (ΔnfnAB) | AG1715 (Δgogat) | Reference strain (LL345) | AVM064 (Δppdk ΔmalE::P<sub>eno</sub><sup>-</sup> pyk) | AVM003 (Δppdk) | LL1084 (ΔnfnAB) | AG1715 (Δgogat) |
| Ethanol | 2.12 ± 0.15 | 3.66 ± 0.11 | 1.99 ± 0.11 | 2.12 ± 0.05 | 1.80 ± 0.14 | 2.65 ± 0.05 | 3.94 ± 0.20 | 2.02 ± 0.13 | 2.46 ± 0.17 | 1.63 ± 0.09 |
| Acetate | 3.03 ± 0.01 | 1.53 ± 0.08 | 3.02 ± 0.04 | 3.04 ± 0.03 | 2.96 ± 0.04 | 2.33 ± 0.01 | 1.54 ± 0.07 | 2.81 ± 0.18 | 2.48 ± 0.09 | 1.77 ± 0.15 |
| Formate | 0.25 ± 0.08 | 0.53 ± 0.04 | 0.20 ± 0.03 | 0.68 ± 0.02 | 0.16 ± 0.02 | 0.37 ± 0.03 | 0.80 ± 0.09 | 0.43 ± 0.11 | 0.31 ± 0.07 | 0.62 ± 0.24 |
| Lactate | 0.03 ± 0.00 | 0.05 ± 0.01 | 0.04 ± 0.00 | 0.03 ± 0.00 | 0.03 ± 0.00 | 0.17 ± 0.01 | 0.10 ± 0.04 | 0.15 ± 0.06 | 0.14 ± 0.02 | 0.08 ± 0.04 |
| Pyruvate | 0.01 ± 0.00 | 0.01 ± 0.00 | 0.03 ± 0.02 | 0.02 ± 0.00 | 0.02 ± 0.00 | 0.22 ± 0.01 | 0.03 ± 0.01 | 0.05 ± 0.01 | 0.08 ± 0.01 | 0.05 ± 0.03 |
| Malate | 0.01 ± 0.01 | 0.26 ± 0.05 | <0.01 | <0.01 | <0.01 | <0.01 | 0.09 ± 0.01 | <0.01 | 0.01 ± 0.00 | <0.01 |
| CO<sub>2</sub><sup>c</sup> | 5.21 ± 0.10 | 4.73 ± 0.19 | 5.14 ± 0.10 | 4.81 ± 0.08 | 4.93 ± 0.09 | 5.15 ± 0.04 | 4.97 ± 0.17 | 4.94 ± 0.18 | 5.16 ± 0.23 | 3.13 ± 0.33 |
| H<sub>2</sub> | 10.6 ± 0.3 | N.D. | N.D. | N.D. | 10.2 ± 0.2 | 6.1 ± 0.3 | N.D. | N.D. | N.D. | 9.0 ± 1.1 |
| Ex. protein<sup>d</sup> | 0.20 ± 0.01 | 0.13 ± 0.01 | 0.16 ± 0.01 | 0.21 ± 0.01 | 0.17 ± 0.00 | 0.12 ± 0.00 | 0.09 ± 0.01 | 0.13 ± 0.00 | 0.11 ± 0.01 | 0.09 ± 0.01 |
| Total amino acids<sup>e</sup> | 0.06 ± 0.00 | 0.07 ± 0.01 | 0.06 ± 0.00 | 0.07 ± 0.00 | 0.05 ± 0.00 | 0.29 ± 0.01 | 0.11 ± 0.02 | 0.26 ± 0.08 | 0.28 ± 0.02 | 0.08 ± 0.02 |
| Unit: μmol g<sub>biomass</sub><sup>-1</sup> h<sup>-1</sup> |  |  |  |  |  |  |  |  |  |  |
| Alanine | 14.2 ± 0.5 | 15.8 ± 1.4 | 13.9 ± 1.1 | 15.0 ± 0.5 | 12.0 ± 0.2 | 23.7 ± 0.3 | 15.6 ± 1.3 | 21.6 ± 1.9 | 19.6 ± 0.9 | 11.3 ± 1.1 |
| Asparagine | 0.3 ± 0.0 | 0.6 ± 0.1 | 0.3 ± 0.1 | <0.2 | <0.2 | <0.2 | <0.2 | <0.2 | <0.2 | <0.2 |
| Aspartate | 0.6 ± 1.1 | 5.2 ± 0.7 | 2.1 ± 0.2 | 2.0 ± 1.0 | 2.3 ± 1.5 | 0.8 ± 0.1 | 2.0 ± 0.5 | 0.6 ± 0.0 | 1.7 ± 0.2 | 0.3 ± 0.2 |
| Cysteine | -80 ± 1 | -304 ± 11 | -324 ± 5 | -335 ± 11 | -73 ± 54 | -262 ± 117 | -409 ± 20 | -404 ± 128 | -473 ± 21 | -66 ± 29 |
| Glycine | 4.5 ± 0.5 | 5.7 ± 0.7 | 4.4 ± 0.3 | 4.1 ± 0.1 | 3.2 ± 0.4 | 5.4 ± 0.1 | 4.1 ± 0.5 | 4.7 ± 0.6 | 4.8 ± 0.4 | 0.8 ± 0.4 |
| Glutamate | 6.1 ± 0.2 | 6.3 ± 1.0 | 4.6 ± 0.1 | 6.9 ± 0.5 | 5.9 ± 0.3 | 14.4 ± 0.3 | 10.5 ± 0.4 | 11.8 ± 0.9 | 20.4 ± 0.7 | 11.6 ± 1.4 |
| Glutamine/arginine<sup>f</sup> | 1.5 ± 0.1 | 4.8 ± 1.1 | 4.3 ± 1.3 | 3.7 ± 0.4 | 1.3 ± 0.1 | 1.2 ± 0.5 | 5.4 ± 0.6 | 6.6 ± 0.8 | 2.4 ± 0.3 | 0.8 ± 0.1 |
| Histidine | <0.2 | <0.2 | <0.2 | <0.2 | <0.2 | <0.2 | 0.5 ± 0.1 | 0.8 ± 0.2 | 0.3 ± 0.0 | <0.2 |
| Isoleucine | 2.8 ± 0.1 | 1.5 ± 0.1 | 3.1 ± 0.1 | 3.6 ± 0.5 | 2.3 ± 0.0 | 24.9 ± 0.7 | 9.3 ± 1.2 | 27.1 ± 7.1 | 22.2 ± 2.0 | 5.2 ± 1.8 |
| Leucine | 1.8 ± 0.0 | 3.4 ± 0.3 | <0.2 | 1.5 ± 1.3 | 1.6 ± 0.0 | 6.2 ± 0.2 | 5.8 ± 0.5 | 7.0 ± 0.4 | 6.9 ± 0.2 | 2.3 ± 0.4 |
| Methionine | <0.2 | <0.2 | <0.2 | <0.2 | <0.2 | 15.6 ± 1.3 | 0.3 ± 0.1 | 1.0 ± 1.2 | 2.2 ± 0.6 | 0.5 ± 0.3 |
| Phenylalanine | 2.4 ± 0.2 | 9.9 ± 1.2 | 2.4 ± 0.5 | 2.0 ± 0.4 | 2.0 ± 0.1 | 10.3 ± 0.2 | 11.1 ± 3.4 | 10.0 ± 1.5 | 14.2 ± 1.1 | 9.8 ± 1.8 |
| Serine | 5.7 ± 0.5 | 4.9 ± 0.5 | 4.7 ± 0.2 | 5.5 ± 0.2 | 4.3 ± 0.2 | 4.8 ± 0.4 | 3.5 ± 0.3 | 4.7 ± 0.1 | 5.0 ± 0.4 | 2.4 ± 0.9 |
| Threonine | 7.3 ± 0.5 | 9.0 ± 1.1 | 7.3 ± 0.7 | 7.8 ± 0.5 | 6.2 ± 0.7 | 9.1 ± 0.3 | 9.2 ± 1.1 | 7.2 ± 1.2 | 12.7 ± 1.0 | 1.0 ± 0.7 |
| Tryptophan | <0.2 | 0.8 ± 0.1 | <0.2 | <0.2 | <0.2 | <0.2 | <0.2 | <0.2 | <0.2 | <0.2 |
| Tyrosine | 2.0 ± 0.1 | 1.9 ± 0.2 | 1.1 ± 0.1 | <0.2 | <0.2 | 4.7 ± 0.2 | 2.9 ± 0.3 | 3.3 ± 0.1 | 9.8 ± 2.7 | <0.2 |
| Valine | 7.2 ± 0.3 | 3.0 ± 0.2 | 10.5 ± 0.2 | 12.7 ± 0.5 | 4.9 ± 0.2 | 167.4 ± 5.1 | 32.1 ± 7.1 | 150.8 ± 63.0 | 159.8 ± 19.1 | 33.9 ± 20.5 |

<sup>a</sup> Data are shown as average ± standard deviation for three biological replicates. <sup>b</sup> Negative values represent consumption and positive values represent production <sup>c</sup> Calculated from fermentation product yields and biomass yield (see Materials and Methods) <sup>d</sup> Based on a molecular weight of a general protein (see Supplemental Table S2 and Materials and Methods). <sup>e</sup> Excluding cysteine. <sup>f</sup> Sum of the two amino acids. N.D. not determined.

**TABLE S7.**
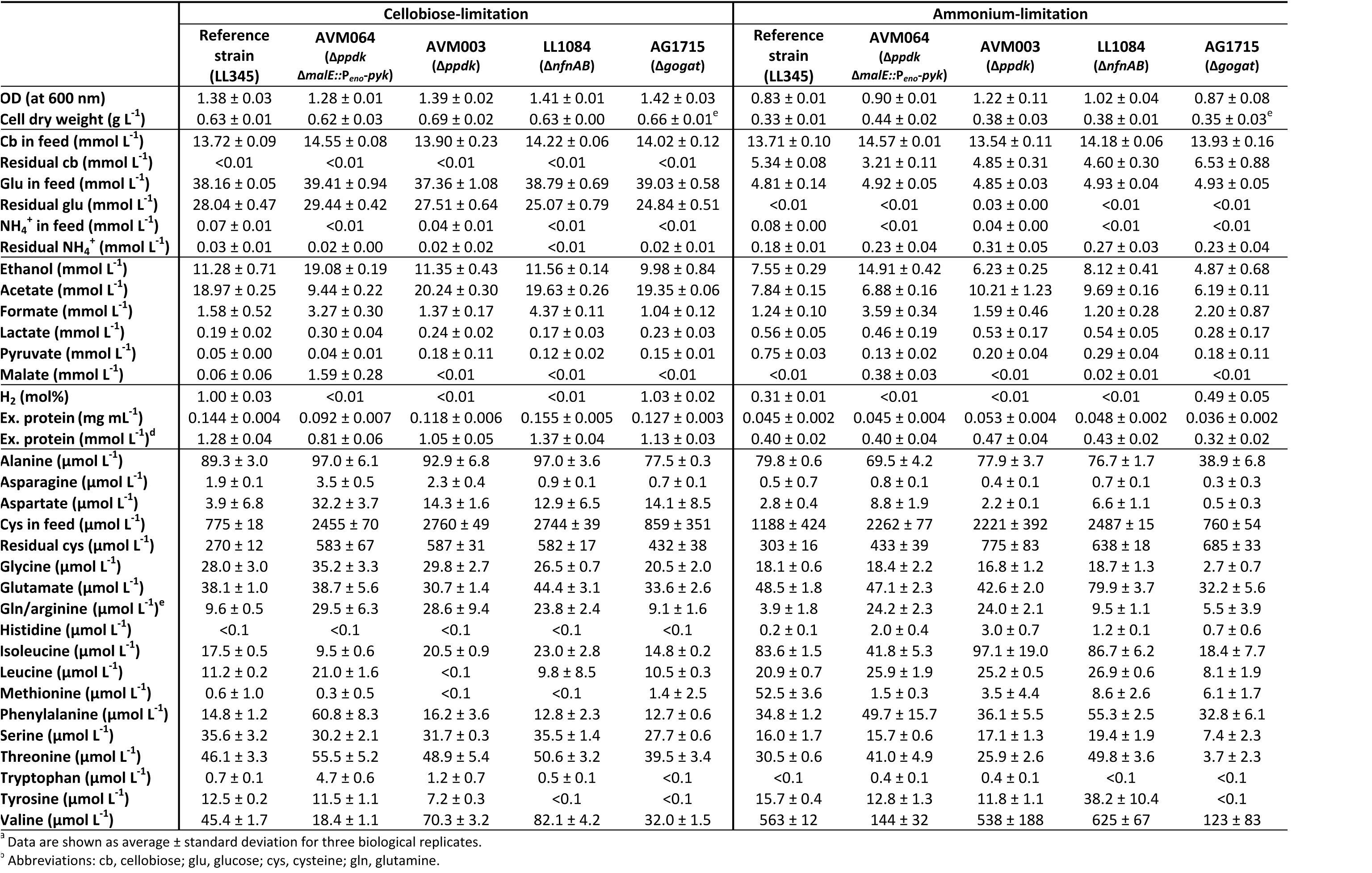
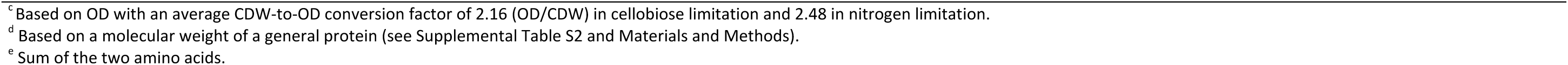
Concentrations of substrates and products in chemostats limited on either cellobiose or ammonium at a dilution rate of 0.10 h^-1^.^a,b^

**TABLE S8.** Lactate dehydrogenase activities for different strains in this study at different conditions, expressed in µmol mg_protein_^-1^ min^-1^.^a,b^

| Strain | Exponential<br>phase in batch | Cellobiose-limitation | Ammonium-limitation |
| --- | --- | --- | --- |
| **WT** | 0.51 ± 0.03 | N.D. | N.D. |
| **AVM003 ($\Delta ppdk$)** | 0.30 ± 0.06 | N.D. | N.D. |
| **AVM064 ($\Delta ppdk \Delta me::P_{\text{eno}}-pyk$)** | 0.33 ± 0.03 | N.D. | N.D. |
| **LL345 ($\Delta hpt$)** | N.D. | 0.69 ± 0.02 | 2.65 ± 0.26 |
| **AG1715 ($\Delta hpt \Delta gogat$)** | N.D. | 0.40 ± 0.02 | 0.9 ± 0.03 |

<sup>a</sup> Data are shown as average ± standard deviation for four technical replicates of biological duplicates.
N.D. not determined

